# Awake hippocampal-prefrontal replay mediates spatial learning and decision making

**DOI:** 10.1101/632042

**Authors:** Justin D. Shin, Wenbo Tang, Shantanu P. Jadhav

## Abstract

Spatial learning requires remembering and choosing paths to goals. Hippocampal place cells replay spatial paths during immobility in reverse and forward order, offering a potential mechanism. However, how replay mediates both goal-directed learning and memory-guided decision making is unclear. We therefore continuously tracked replay in the same hippocampal-prefrontal ensembles throughout learning of a spatial alternation task. We found that during pauses between behavioral trajectories, awake reverse and forward hippocampal replay consistently mediated an internal cognitive search of all available past and future possibilities, and exhibited opposing learning gradients for prediction of past and future behavioral paths, respectively. Coordinated hippocampal-prefrontal replay mediated recall of correct past paths and selection of future choices leading to reward based on the hippocampal cognitive search, executing spatial working memory rules. Our findings reveal a learning shift from hippocampal reverse-replay-based retrospective evaluation to forward-replay-based prospective planning, with prefrontal filtering of memory-guided paths for learning and decision-making.

## INTRODUCTION

The hippocampus is necessary for formation and retrieval of episodic memories to guide daily behavior, including goal-directed spatial learning and navigation (Eichenbaum and Cohen, 2004; Squire, 1992). Learning spatial paths and choices that lead to desired goals is critical for foraging and survival, and requires animals to remember and plan routes to rewards at distant locations. Hippocampal place cells are active in specific spatial locations during active exploration (O’Keefe and Nadel, 1978). While this spatial code provides information about current location, spatial memories require learning links between sequences of locations that encode specific paths, and choices that lead to goals. Interestingly, hippocampal activity exhibits another phenomenon called “replay”, which is associated with high frequency sharp-wave ripple (SWR) events prevalent during offline periods in sleep and non-exploratory waking states (‘awake replay’) (Buzsáki, 2015; Carr et al., 2011; Foster, 2017; Leonard et al., 2015). During replay, temporally compressed sequences of place cells reactivate spatial trajectories that span parts of the entire environment in either forward or reverse order (Ambrose et al., 2016; Csicsvari et al., 2007; Davidson et al., 2009; Diba and Buzsáki, 2007; Farooq and Dragoi, 2019; Foster and Wilson, 2006; Gupta et al., 2010; Ólafsdóttir et al., 2017; Pfeiffer and Foster, 2013; Tang et al., 2017; Xu et al., 2019). Awake hippocampal replay, seen prominently during pauses in exploration and consummatory behavior, is known to be necessary for spatial learning, especially for spatial working memory tasks (Jadhav et al., 2012), and has been proposed to potentially provide neural correlates of various memory processes, notably memory consolidation, recall, and decision making (Buzsáki, 2015; Carr et al., 2011; Foster, 2017; Joo and Frank, 2018; Tang and Jadhav, 2018).

Evidence from several studies supports a role of awake replay in spatial memory-guided behavior (Ambrose et al., 2016; Diba and Buzsáki, 2007; Dupret et al., 2010; Foster and Wilson, 2006; Gupta et al., 2010; Ólafsdóttir et al., 2017; Papale et al., 2016; Pfeiffer and Foster, 2013; Singer et al., 2013; Tang et al., 2017; Vaz et al., 2019; Wu et al., 2017; Xu et al., 2019). Reverse replay is enhanced by reward and novelty (Ambrose et al., 2016; Cheng and Frank, 2008; Foster and Wilson, 2006; Singer et al., 2013), leading to suggestions that it could support temporal credit assignment for associating sequential spatial experiences that encode paths with resultant outcomes (Ambrose et al., 2016; Carr et al., 2011; Foster, 2017; Foster and Knierim, 2012; Haga and Fukai, 2018; Mattar and Daw, 2018; Pfeiffer, 2018). On the other hand, replay has also been proposed to support memory retrieval for planning upcoming behaviors (Carr et al., 2011; Pfeiffer and Foster, 2013; Singer et al., 2013; Wu et al., 2017), with a potential role of forward replay in planning (Xu et al., 2019). Despite this evidence and proposed roles (Buzsáki, 2015; Carr et al., 2011; Foster and Knierim, 2012; Haga and Fukai, 2018; Mattar and Daw, 2018; Pezzulo et al., 2014; Tang and Jadhav, 2018), how reverse and forward replay mediate both novel spatial learning and recall for decision making remains unclear, since this requires monitoring the evolution of content of replay events in the same neural populations over the entire duration of learning a replay-dependent task with memory-guided choices toward goals.

Furthermore, goal-directed learning and planning relies on a wider neural network including not only the hippocampus, but also regions involved in the evaluation and selection of task-relevant memories during retrieval and decision making, notably the prefrontal cortex (PFC) (Benchenane et al., 2011; Eichenbaum, 2017; Pezzulo et al., 2014; Redish, 2016; Spellman et al., 2015; Tang and Jadhav, 2018; Yu and Frank, 2015). The cognitive processes of learning and deliberation are known to require the PFC (Yu and Frank, 2015), and PFC involvement in navigation and spatial memory is established in both humans and rodents (Eichenbaum, 2017; Epstein et al., 2017). How hippocampal and prefrontal networks together support novel learning and planning, especially for replay-dependent working memory tasks, remains an open question (Eichenbaum, 2017; Pezzulo et al., 2014; Tang and Jadhav, 2018). SWRs have been shown to mediate coordinated hippocampal-prefrontal reactivation of spatial paths (Jadhav et al., 2016; Peyrache et al., 2009; Tang et al., 2017), but whether and how this coordinated reactivation plays a role in memory-guided behavior is not known.

## RESULTS

### Continuous tracking of forward and reverse replay throughout learning

In order to address these questions, we used continuous and simultaneous electrophysiological monitoring of ensembles of hippocampal-prefrontal neurons in rats learning a novel, replay-dependent W-track spatial alternation task in a single day (**Figure 1**; **Figures S1** and **S2**) (Jadhav et al., 2012; Maharjan et al., 2018). This task involves continuous alternation between reward wells on the three arms of the W-track (**Figure 1A**), and animals are rewarded upon completion of a correct inbound or outbound behavioral sequence according to the following rules: (i) starting from either side well, animals have to return to the center well (inbound trajectories 2 and 4), and (ii) starting from the center well, animals have to recall the previous inbound trajectory and choose the opposite side well from the previously visited side well (outbound trajectories 1 and 3). Animals therefore have to learn the sequence of reward-well visits, and choose associated paths leading from one well to another (trajectories 1-4, **Figure 1A**) to obtain reward.

**Figure 1.**
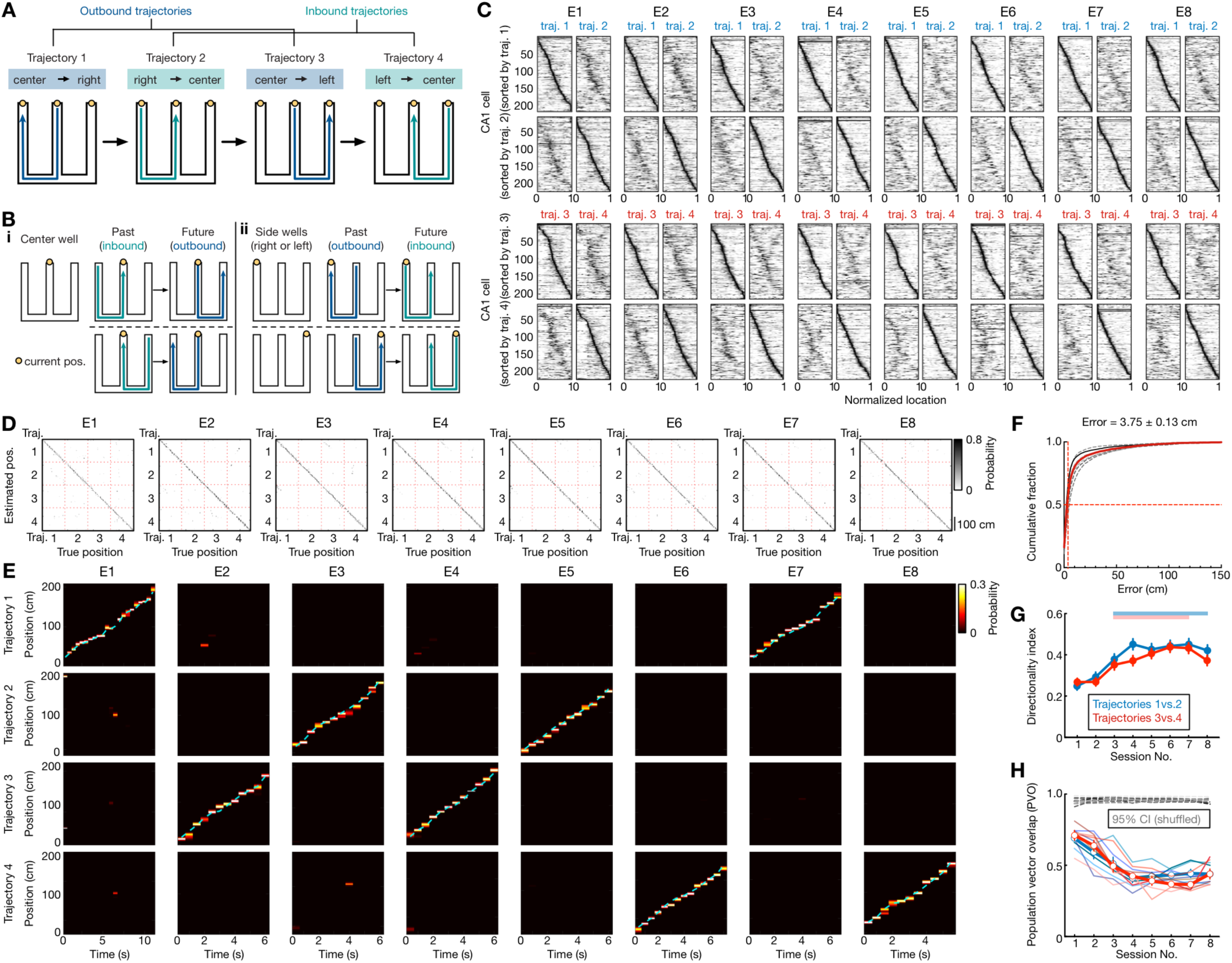
Hippocampal place-cell sequences distinctly code different behavioral trajectories during learning of a W-maze spatial memory task. **(A)** W-maze spatial alternation task design, depicting the correct behavioral sequence of outbound (blue) and inbound (teal) trajectories (labelled 1-4) for reward. Right-left labelled according to animal direction. **(B)** Past and future trajectories during transitions at reward wells in the W-maze task. **(i)** Two possible correct behavioral sequences at the center well, with inbound and outbound trajectories representing possible past and future paths, respectively. A behavioral sequence of inbound past path and outbound future path to the opposite side-arm comprises a history-dependent spatial working memory rule. **(ii)** One correct behavioral sequence for each side well (left or right well). Note that compared to the center well, the past and future paths at side wells are reversed as outbound and inbound trajectories, respectively. The inbound return-to-center future path originating at the side well comprises a history-independent spatial memory rule. **(C)** Place fields of all CA1 place cells (*n* = 216) recorded from 6 rats continuously across 8 learning sessions (or epochs; denoted as E1-8). The fields for trajectories 1 and 2, sorted according to peak positions on trajectory 1 or 2, are shown on the top two rows. The bottom two rows are for trajectories 3 and 4. **(D-F)** Position reconstruction based on CA1 ensemble spiking during active running behavior on trajectories (> 5 cm/s). **(D)** Example confusion matrices (estimated vs. true position) for all place-cell reconstruction. Values along diagonal show correspondence between estimated and true position. **(E)** illustrative estimated position probabilities based on Bayesian decoding of spike trains during running on different trajectories, using data from a single animal for individual sessions. Cyan line: actual animal trajectory. **(F)** Cumulative CA1 decoding errors across all animals (*n* = 6 rats). Dashed lines: individual animals; Red line: all animals; Solid black line: the example animal shown in **(D)**. Median error of all sessions noted on top as (median ± SEM). **(G and H)** Directional selectivity of place cells across different learning sessions. **(G)** Directionality index (DI) of place cells across learning sessions. Note the significant overall increase of DIs in the population from the first to the last sessions (red and blue bars above indicate significant difference from the first session, *p* < 0.05, Friedman tests with Dunn’s *post hoc*). Error bars: SEM. **(H)** Similarity of the place-cell population in two running directions was computed using the population vector overlap (PVO). Dashed grey lines: 95% CIs of the shuffled distributions (by randomly permuting running directions) from each animal. Thin blue and red lines: actual PVO from individual animals on right (i.e., trajectories 1 vs. 2) and left (i.e., trajectories 3 vs. 4) trajectories, respectively. Thick lines and error bars: means and SEM. Note that the similarity of the place-field templates in two running directions decreased across sessions as expected from the increase in the DIs, but distinct templates for each direction are apparent as early as the first session (*p’s* < 0.0001 compared to the shuffled data for individual rats, permutation tests) (Foster, 2017; Foster and Wilson, 2006). See also **Figures S1** and **S2**, and **Table S1**.

Both awake replay and functional hippocampal-prefrontal interactions are necessary for learning the outbound component of this task, which requires spatial working memory (Jadhav et al., 2012; Maharjan et al., 2018). A correct spatial working memory behavioral sequence consists of two consecutive trajectories with a transition at the center well (**Figure 1B_i_**), where the past path is an inbound trajectory that terminates at the center well, and the future path is an outbound trajectory that originates at the center well and proceeds to the opposite side-arm. The outbound, spatial working memory component is therefore history-dependent (Jadhav et al., 2012). In contrast, the inbound component simply requires implementation of a “return-to-center” rule from each side-well (**Figure 1B_ii_**), and this spatial reference memory rule is history-independent. For side well transitions, the past path in a correct behavioral sequence is an outbound trajectory terminating at the side well, and the future path is an inbound trajectory originating at the side well. The past and future paths are thus reversed at the center and side wells (**Figure 1B**).

Animals were tasked with learning the W-maze rules in eight behavioral sessions (epochs 1-8, denoted as E1-8, 15-20 mins per session) in a single experimental day, interleaved with rest sessions in a sleep box (**Figure S2**; see **STAR Methods**) (Maharjan et al., 2018; Tang et al., 2017). We used custom-built 32 tetrode microdrive arrays to continuously and simultaneously record activity from dorsal CA1 place cells and medial prefrontal cortex (PFC, pre-limbic and anterior cingulate cortical regions) in six rats over the course of learning the task (see **STAR Methods**; recording locations in all animals indicated in **Figures S1A-S1C**; number of cells recorded in each animal shown in **Figures S1D-S1G** and **Table S1**). Activity was monitored with the high temporal resolution of electrophysiological recordings from the same stable CA1 (*n* = 216) and PFC (*n* = 154) ensembles throughout the duration of the entire experiment (5.5-6.5 hours; **Figures S1D-S1G** shows isolation and stability parameters for all neurons recorded). All six animals exhibited rapid learning over the course of the 8 behavioral sessions (learning curves shown in **Figure S2**, see **STAR Methods**), and the same CA1-PFC ensembles were continuously monitored for each animal over the 8 learning sessions. This experimental design thus enabled investigation of CA1 and coherent CA1-PFC replay dynamics using the same ensembles, starting from initial acquisition of the task and through later performance at above-chance levels.

CA1 place cells exhibited spatial and direction selectivity, and formed unique sequential representations of different trajectories during learning (**Figures 1C-1H**). **Figure 1C** shows responses of all recorded CA1 place cells for the 4 trajectories of the task, and for each of the 8 behavioral sessions (sorted by order of place field peaks). Notably, place cell encoding of spatial locations enabled highly accurate decoding of animal position during running on trajectories across all sessions (**Figures 1D-1F**). Second, comparison of the respective outbound-inbound trajectory pairs (1 vs. 2, and 3 vs. 4 in **Figure 1A**) confirmed previous reports that CA1 place cells were directionally selective starting with the first session on the novel track (Foster and Wilson, 2006), and direction selectivity significantly improved over experience from sessions 1-8 (**Figures 1G** and **1H**) (Navratilova et al., 2012; Xu et al., 2019). Place-field templates of the 4 trajectories were thus significantly distinguishable for all epochs, despite a small fraction of bidirectional cells (26.8 ± 12.4%, mean ± SD; see **Table S1** for the number of bidirectional cells in each animal).

For each animal, the 4 behavioral trajectories thus had unique template place-cell sequence representations from stably recorded ensembles starting from the first behavior session, enabling unambiguous detection and continuous tracking of forward and reverse hippocampal replay events (**Figures 1C-1H**). To investigate the content of replay over learning, we used established methods to detect SWRs and candidate replay events during immobility periods at reward wells, and used Bayesian decoding to identify CA1 sequential replay events, with each event distinctly determined as forward or reverse replay of one of the four trajectories (see **STAR Methods**). Examples of forward and reverse replay sequences, corresponding to different stages in learning and detected from the same hippocampal population in one animal, are shown in **Figures 2A-2F** (additional examples in **Figure S3**). Each example shows place-cell activity during a SWR candidate event, templates for all 4 trajectories (linearized place fields of cells sorted by their peak locations on the detected replay trajectory), and Bayesian reconstruction of the animal’s trajectory, with the strength of replay and statistical significance based on time-shuffled data indicated. During immobility periods at reward-well transitions between trajectories where the animals stopped in order to receive reward (immobility time: 10.3 ± 5.7 sec in mean ± SD; **Figure 2G**), multiple SWRs and replay candidate events were prevalent (**Figures 2H** and **2I;** immobility periods with ≥ 2 events, 85.7%, 1313 out of 1533 trials for SWRs; 53.4%, 818 out 1533 trials for replay candidate events). Further, there was no overall bias toward reverse or forward replay of any particular trajectory type (**Figures 2J** and **2K**).

**Figure 2.**
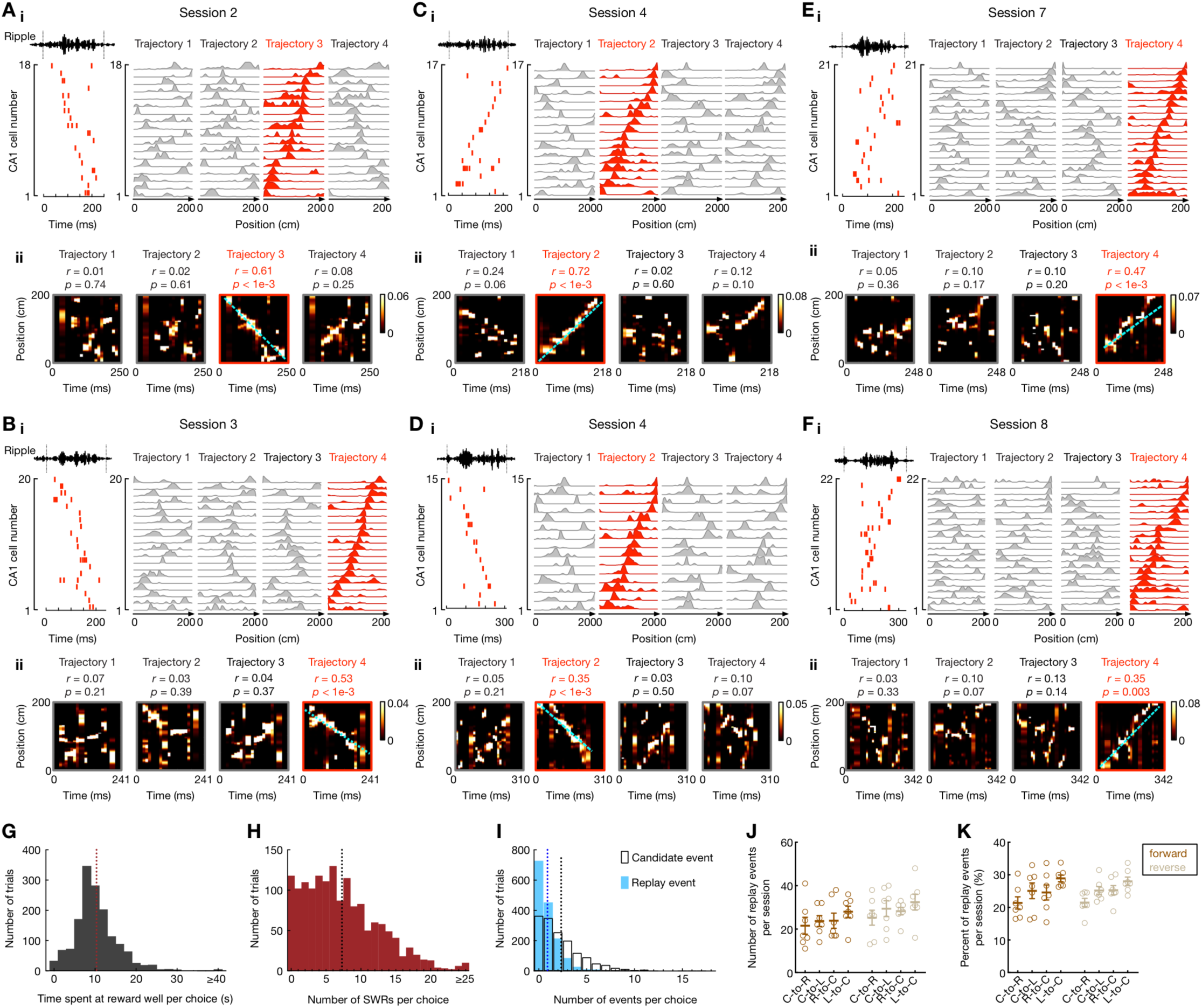
Continuous tracking of forward and reverse replay throughout learning. **(A-F)** Six examples of forward and reverse replay of behavioral trajectories in different learning sessions using continuously tracked CA1 ensembles in one animal. **(i)** *Left*: Place-cell activity during a SWR, with ripple-filtered LFP (150-250 Hz) from one tetrode shown on top (black line). *Right*: Corresponding linearized place fields on trajectories 1 to 4 sorted by their peak locations on the replay trajectory (red). **(ii)** Bayesian reconstruction of the decoded behavioral trajectories with the replay quality (*r*) and *p*-value based on time-shuffled data denoted on top. Cyan lines: the linear fit maximizing the likelihood along the replay trajectory. Color bars: posterior probability. See also **Figure S3** for additional replay examples. **(G-I)** Distributions of **(G)** animals’ immobility times at reward wells, **(H)** the number of SWRs, and **(I)** candidate events (open bars) and replay events (solid bars) detected in immobility periods per choice (i.e., per transition). Only correct trials are shown. The vertical dashed lines on the histograms represent the mean values (Immobility time: 10.3 ± 5.7 sec; SWRs: 7.3 ± 5.5 events; Candidate events: 2.3 ± 2.4 events; replay events: 0.9 ± 1.2 events; data are presented in mean ± SD). **(J** and **K) (J)** Number and **(K)** percentage of forward and reverse replay events of different trajectory types. Each dot represents a session, summing over all 6 animals (sessions 1 and 2 were combined). Error bars: mean ± SEM. Note that the number of replay events was similar across different trajectory types (*F*(7, 42) = 3.062 and *p* = 0.053 for **J**, *F*(7, 42) = 2.227 and *p* = 0.12 for **K**, respectively; repeated measures ANOVA).

### Reverse replay of possible past paths, forward replay of available future paths

In order to examine the relationship between replay content and behavioral choices, we focused our analyses on immobility periods during the trajectory switching phase at reward wells. Examples of correct behavioral sequences comprising two consecutive trajectories are illustrated for transitions at a side well (**Figures 3A-3C**) and the center well (**Figures 3D-3F**), respectively. The side-well transition consists of an outbound past trajectory (RUN1, center-to-left), followed by an inbound future trajectory (RUN2, left-to-center), with the unique directional place-cell trajectory sequences for RUN1 and RUN2 depicted in the sorted CA1 place-cell activity (**Figure 3A**). Three replay events occurred during this transition at the side reward well, identified as two reverse replay events of the past trajectory (RUN1), and one forward replay event of the future trajectory (RUN2) (**Figure 3B**; templates for all trajectories 1-4 in **Figure 3C**), in an inter-mixed order of occurrence (reverse-forward-reverse). The center-well transition example consists of an inbound past trajectory (RUN1, left-to-center), followed by an outbound future trajectory (RUN2, center-to-right) (**Figure 3D**). Place-cell trajectory sequences for RUN2 and the alternative inbound past trajectory (RUN1-alt, right-to-center) are depicted in the sorted CA1 place cell activity (**Figure 3D**; templates for all trajectories 1-4 in **Figure 3F**). Interestingly, the four detected replay events at the center well comprised of three forward replay events of the future taken path, and one reverse replay event of the *alternative* inbound past path, not the actual past path taken by the animal (**Figure 3E**; note that at the center well, the two inbound trajectories represent the two possible past paths, and the two outbound trajectories represent the two possible future paths, with past and future paths reversed at the side wells). The behaviorally actualized path by the animal is denoted as the ‘taken’ path, and the alternative path is denoted as the ‘not-taken’ path. **Figures 3G-3I** show three additional examples for side and center wells, with decoded replay paths depicted on the track (see **STAR Methods**; further examples during different learning stages in **Figure S4**). Replay events in the side well example (**Figure 3G**) again reactivate the animal’s taken past and future path in the reverse and forward order, respectively, whereas at the center well (**Figures 3H** and **3I**), the replay events reactivate the alternative (not-taken) future (**Figure 3H**) and alternative (not-taken) past (**Figure 3I**) paths, but still maintain the forward and reverse order, respectively.

**Figure 3.**
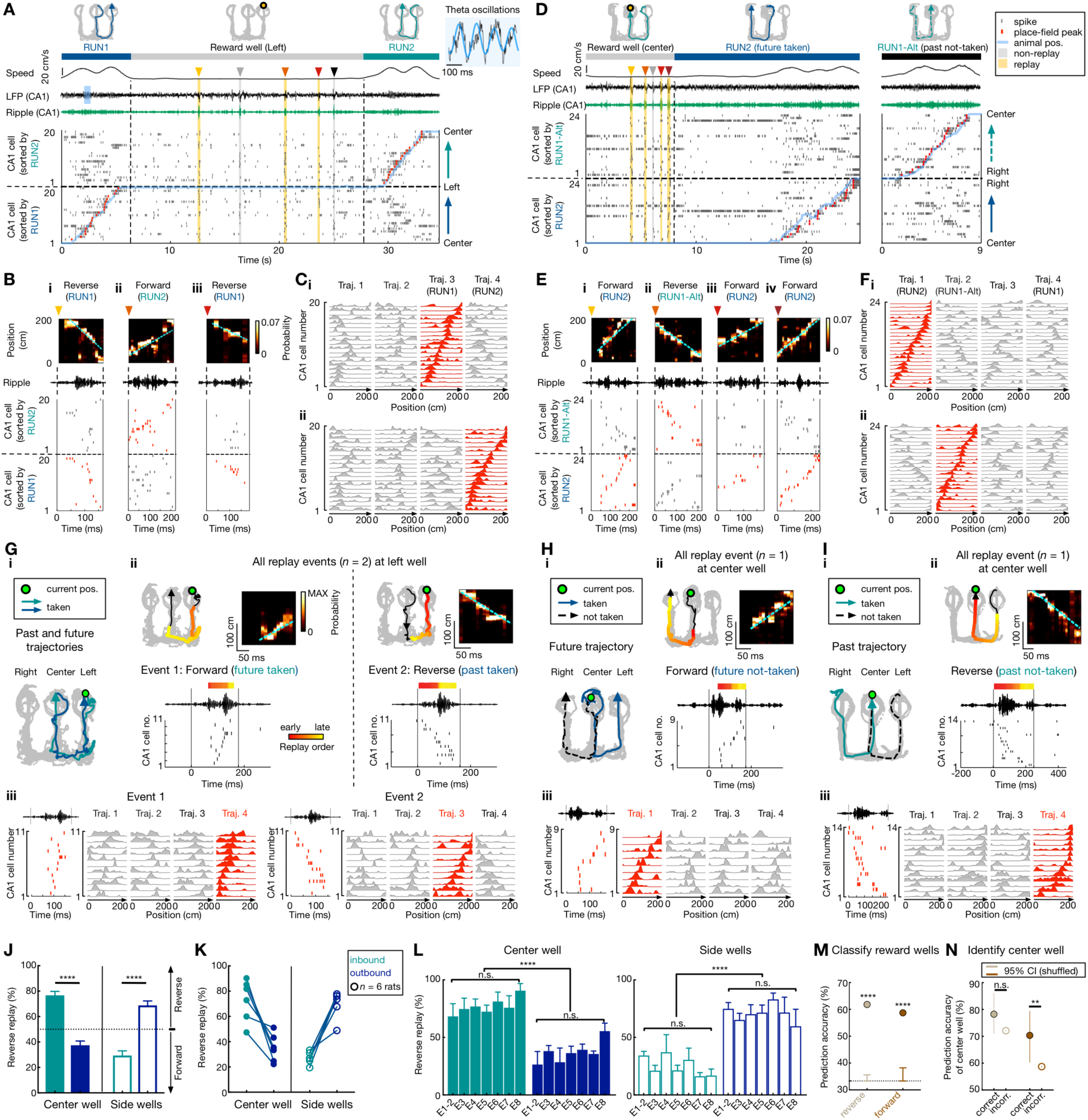
Reverse replay of possible past paths, forward replay of possible future paths. **(A-F)** Illustration of past and future trajectories replayed by reverse and forward place-cell sequences at reward wells. **(A-C)** Example of side reward well transitions. **(A)** CA1 neural activity during a correct outbound-inbound sequence (RUN1 to RUN2), with replay events at the left side well. From *top* to *bottom*, the behavioral sequence (blue and teal lines represent the past and future taken paths, RUN1 and RUN2; grey segment denotes the immobility period at reward well), animal speed, broadband and ripple-filtered LFPs from one CA1 tetrode, raster plot of 20 place cells ordered by the positions of their place-field peaks (red ticks; overlaid blue line indicates linearized position of the animal). CA1 exhibits theta oscillatory activity (8-10 Hz; blue shading on LFP) during running, and large amplitude SWRs during immobility at the reward well, coincident with synchronous activity of place cells (indicated by arrowheads and shadings in the raster plot). Five synchronous events were seen, with 3 events detected as significant replay sequences (yellow shading). **(B)** Expanded view of the 3 replay events from **(A)**, identified as forward replay of RUN2 (future) and reverse replay of RUN1 (past). **(B_i_-B_iii_)** Each column shows the Bayesian reconstruction of the animal’s trajectory during replay (*top*), ripple-filtered LFP (*middle*), and spike raster of place cells sorted as in **(A)** (*bottom*; raster corresponding to the replay trajectory is in red). **(C)** Linearized fields of the place cells in **(A)** for trajectories 1 to 4 sorted by their peak locations on **(C_i_)** RUN1 and **(C_ii_)** RUN2. **(D-F)** Example of center reward well transitions. **(D)** CA1 neural activity during a correct inbound-outbound sequence (RUN1 to RUN2), with replay events at the center well. RUN1-Alt: alternative past trajectory to center well. **(E)** All four replay events during the center-well transition, identified as forward replay of RUN2 (future) and reverse replay of RUN1-Alt (*alternative* past). **(F)** Linearized fields of the place cells in **(D)**. Data are presented as in **(A-C)**. **(G)** Replay events during an example transition at the left side well. **(i)** Behavioral sequences for the side-well transition. **(ii)** All replay events (*n* = 2; Event 1 and Event 2) during the transition at the side well. *Top left:* Bayesian reconstruction of the replay trajectory. Colored points overlaid on the replay trajectory (black arrowhead line) indicate the Bayesian-decoded positions, with the color denoting relative time within the replay event (early in red, and late in yellow). *Top right*: Bayesian reconstruction of the decoded behavioral trajectories. Cyan lines: the linear fit maximizing the likelihood along the replay trajectory. *Bottom*: ripple-filtered LFP and the sequential spiking of place cells during the SWR. **(iii)** Linearized fields of the place cells active during the Event 1 (*left*) and Event 2 (*right*) on trajectories 1 to 4 sorted by their peak locations on the replay trajectory (red). **(H** and **I)** Replay events during two example transitions at the center well. Data are presented as in **(G)**. See also **Figure S4** for additional replay examples. **(J)** At the center well, inbound trajectories (past paths) and outbound trajectories (future paths) were preferentially replayed in a reverse and forward order, respectively. At the side wells, outbound trajectories (past) and inbound trajectories (future) were preferentially replayed in a reverse and forward order, respectively. \*\*\*\**p* < 1e-4, session-by-session rank-sum paired test. Error bars: SEM. **(K)** The bias in **(J)** is consistent in each rat (circles; *n* = 6). **(L)** The bias in **(J)** at the center (*left*) and side (*right*) wells appeared similarly across 8 behavioral sessions (denoted as E1-8). *n.s*.: non-significant (Center well: main effect of group, *p* = 0.52 and 0.44 for inbound and outbound, respectively; Side wells: main effect of group, *p* = 0.30 and 0.59 for inbound and outbound, respectively; Kruskal-Wallis tests). \*\*\*\**p* < 1e-4, session-by-session rank-sum paired test. **(M)** Patterns of forward and reverse replay discriminate different goal-locations (*n* = 3 wells). The prediction accuracy of classifying wells is shown based on the number of either forward or reverse replay events for different trajectories at the wells. The cross-validated decoder using SVM is significantly better than chance (error bars) defined by permutation tests (\*\*\*\**p* < 0.0001). **(N)** The prediction accuracy of incorrect trials is significantly lower than that of correct trials for forward replay events, but not reverse replay events at the center well. Error bars indicate 95% confidence intervals based on bootstrapped data (Correct trials were randomly subsampled 1,000 times to match the number of incorrect trials; *n.s*., *p* = 0.06; \*\**p* = 0.01). See also **Figures S4 and S5**.

We quantified this relationship of forward and reverse replay content at center and side reward wells to ongoing behavioral trajectory sequences during correct trials. Similar to examples in **Figure 3**, there was a strong and consistent prevalence of reverse replay of the two possible past choices (actual taken and alternative past paths *to* reward well), and forward replay of the two possible future choices (actual taken and alternative future paths *from* reward well), associated with the respective reward well location (**Figures 3J** and **3K**; **Figure S5**). At the center well, this manifested as reverse replay of inbound trajectories (possible past paths; reverse/forward events, 324/91, *p* < 1e-4, z-test for proportions), and forward replay of outbound trajectories (possible future paths; reverse/forward events, 116/202, *p* < 1e-4, z-test for proportions), which was reversed at the side wells (reverse/forward events for outbound trajectories: 267/115, reverse/forward events for inbound trajectories: 101/272, *ps* < 1e-4, z-tests for proportions; session-by-session comparison in **Figure 3J**). This effect was consistent across all six animals (**Figure 3K**), and persisted across all behavioral sessions during different stages in learning (**Figure 3L**).

Reverse replay of past paths and forward replay of future paths persisted when we included only significantly unidirectional CA1 cells to rule out any unintended bias due to bidirectional cells (**Figure S5A**). We also examined if there was a tendency for reverse and forward replay to occur at the beginning and end of immobility periods, respectively (i.e., at the end of previous trajectory and just prior to the upcoming trajectory respectively), as previously reported on linear tracks (Diba and Buzsáki, 2007). No such bias was apparent, similar to recent reports (Ambrose et al., 2016), and reverse replay of past paths and forward replay of future paths continued in an inter-mixed order throughout the immobility periods (**Figures S5B** and **S5C**). We did, however, find that replay rate was significantly higher during ‘disengaged’ periods during the middle periods of reward-well transitions, compared to ‘engaged’ periods near the beginning and end of reward-well transitions (Ólafsdóttir et al., 2017) (**Figures S5D-S5F**). Finally, we also confirmed that this effect was not a result of bias in the distribution of place fields or decoded replay positions (**Figures S5G-S5I**). In fact, the distribution of decoded positions during replay events again revealed the over-representation of past path positions in reverse replay and future path positions in forward replay (**Figures S5H** and **S5I**).

The behavioral relevance of this replay content structure was further confirmed by using the identity of the reverse and forward replay events to predict the current reward well location (i.e. left, center, or right; see **STAR Methods**), with prediction accuracies that were significantly higher than chance-level (**Figure 3M**). Further, we compared this effect of reverse past replay and forward future replay for correct and error working-memory trials that originated from the center well, and observed an impairment specifically in forward replay of future paths. There was a significant decrease in accuracy of predicting the center well using forward replay during error trials compared with correct trials (**Figure 3N**), since the bias of forward replay for future paths was absent in error trials (for error trials, forward replay of future: 45%, *p* = 0.53; reverse replay of past: 68.5%, *p* = 0.008; z-test for proportions).

### Contrasting evolution of reverse and forward replay with learning

Since reverse and forward replay events consisted of both actual taken and alternative (not-taken) past and future paths respectively (**Figure 3**; **Figure S4**), we asked if there was any relationship between the behavioral content of replay events and the memory demands at different stages of learning for the individual task components. We found that even as behavioral performance improved over learning sessions for all animals (**Figure 4A**), there was no change in the balance of overall reverse and forward replay events over the course of learning (**Figures 4B-4H**). SWR rate did not change (**Figure 4B**), but SWR duration and place cell firing during SWRs significantly decreased over the course of learning (**Figures 4C** and **4D**). This corresponded to a decrease in replay rate (**Figure 4E**), although decoded replay length increased with learning (**Figure 4F**). In addition to the decrease in replay rate, as animals became increasingly proficient in the task, there was a decrease in immobility periods at reward wells, resulting in an overall reduction in number of SWRs and replay events at reward wells (session-by-session distributions in **Figure S6**). However, fraction of reverse and forward replay events of any trajectory type did not change, maintaining the balance of reverse and forward replay throughout the course of learning (**Figures 4G** and **4H**).

**Figure 4.**
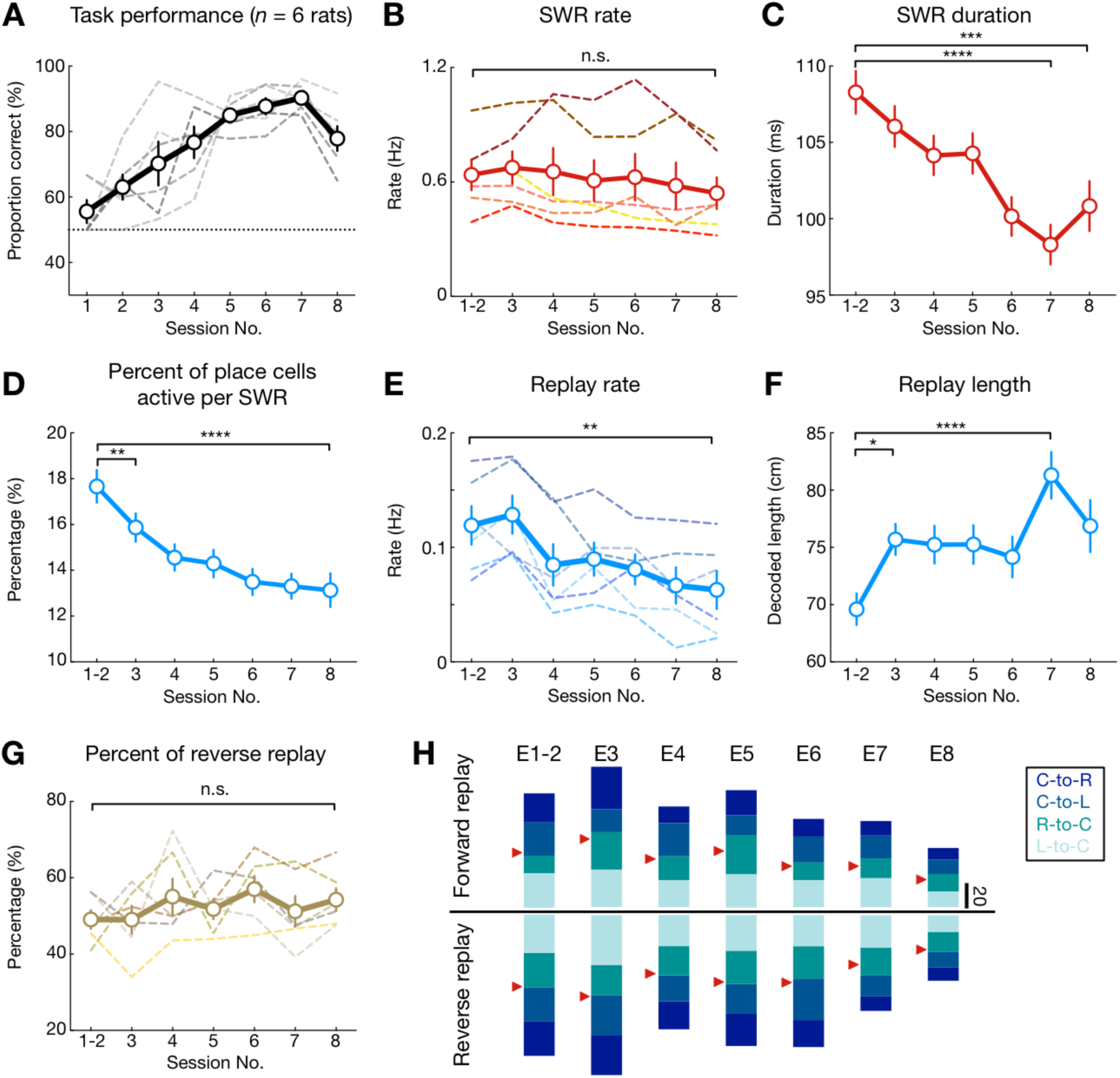
Dynamics of SWR replay properties over the course of learning. **(A)** Task performance, showing proportion correct of outbound trajectories per session for all animals (*n* = 6 rats). **(B)** Total SWR rate did not change across sessions (*p* = 0. 28 and *F*(6, 30) = 1.44, repeated measures ANOVA). **(C)** Duration of SWR events decreased over learning (Kruskal-Wallis test, *p* < 1e-4; \*\*\**p* = 0.0003, \*\*\*\**p* < 1e-4, Dunn’s *post hoc*). Error bars: SEM. **(D)** Percentage of place cells active per SWR was significantly decreased across sessions (\*\*\*\**p* < 0.0001, Kruskal-Wallis test; \*\**p* = 0.0016, \*\*\*\**p* < 0.0001, Dunn’s *post hoc* tests). **(E)** Replay rate decreased over learning (*p* = 0.0005, *F*(6, 30) = 14.21, repeated measures ANOVA; \*\**p* = 0.003, Tukey’s *post hoc* tests). **(F)** Decoded length of replay trajectories increased over learning (Kruskal-Wallis test, *p* = 0.0004; \*\*\**p* = 0.0001, \**p* = 0.039, Dunn’s *post hoc*). Error bars: SEM. **(G)** Percentage of reverse replay out of all replay events did not change across sessions (*p* = 0.12, *F*(6, 30) = 2.370, repeated measures ANOVA). Dashed lines in **(A, B, E, G)**: individual animals. Solid line and error bars: mean and SEMs. **(H)** Number of forward and reverse replay events of the 4 behavioral trajectory types across 8 learning sessions (E1-8) (i.e., center-to-right, C-to-R; center-to-left, C-to-L; right-to-center, R-to-C; left-to-center, L-to-C). The trajectory types are color-coded with red arrowheads indicating 50% level for forward and reverse replay. Note that the fraction of reverse and forward replay events of any trajectory types remained similar across 8 learning sessions. See also **Figure S6**.

We examined replay content at different learning stages independently at center- and side-well transitions, since outbound (center-to-side) and inbound (side-to-center) trajectories *originating* from these reward wells entail distinct memory demands (history-dependent spatial working memory demands and history-independent spatial reference rule, respectively) (Jadhav et al., 2012). Examples of replay events during early learning in initial sessions (E1-3), and late performance in final sessions (E6-8), are shown in **Figures 5A-5D** and **Figure S4** (behavioral performance on the outbound, spatial working memory component, 59.9 ± 9.1% for early or novel learning, and 83.8 ± 9.6% for late performance in mean ± SD; behavioral performance on the inbound memory component, 65.53 ± 28.16% for early or novel learning, and 97.14 ± 0.03% for late performance; see **STAR Methods** and **Figure S2** for all behavior). During early learning, corresponding to low performance levels, reverse replay events at the side wells preferentially reactivated the actual taken past path of the animal, with this bias lost over the course of learning (**Figures 5A** and **5E**); whereas forward replay events at side wells shifted their content from no initial bias during early learning to preferential replay of the future taken path during late sessions, when animals started performing well above chance levels (**Figures 5C** and **5E**). In contrast, at the center well, reverse and forward replay events continued to persistently reactivate, in an unbiased manner, both the actual taken and the alternative (not-taken) past and future paths respectively, throughout the course of learning (**Figures 5B, 5D**, and **5F**).

**Figure 5.**
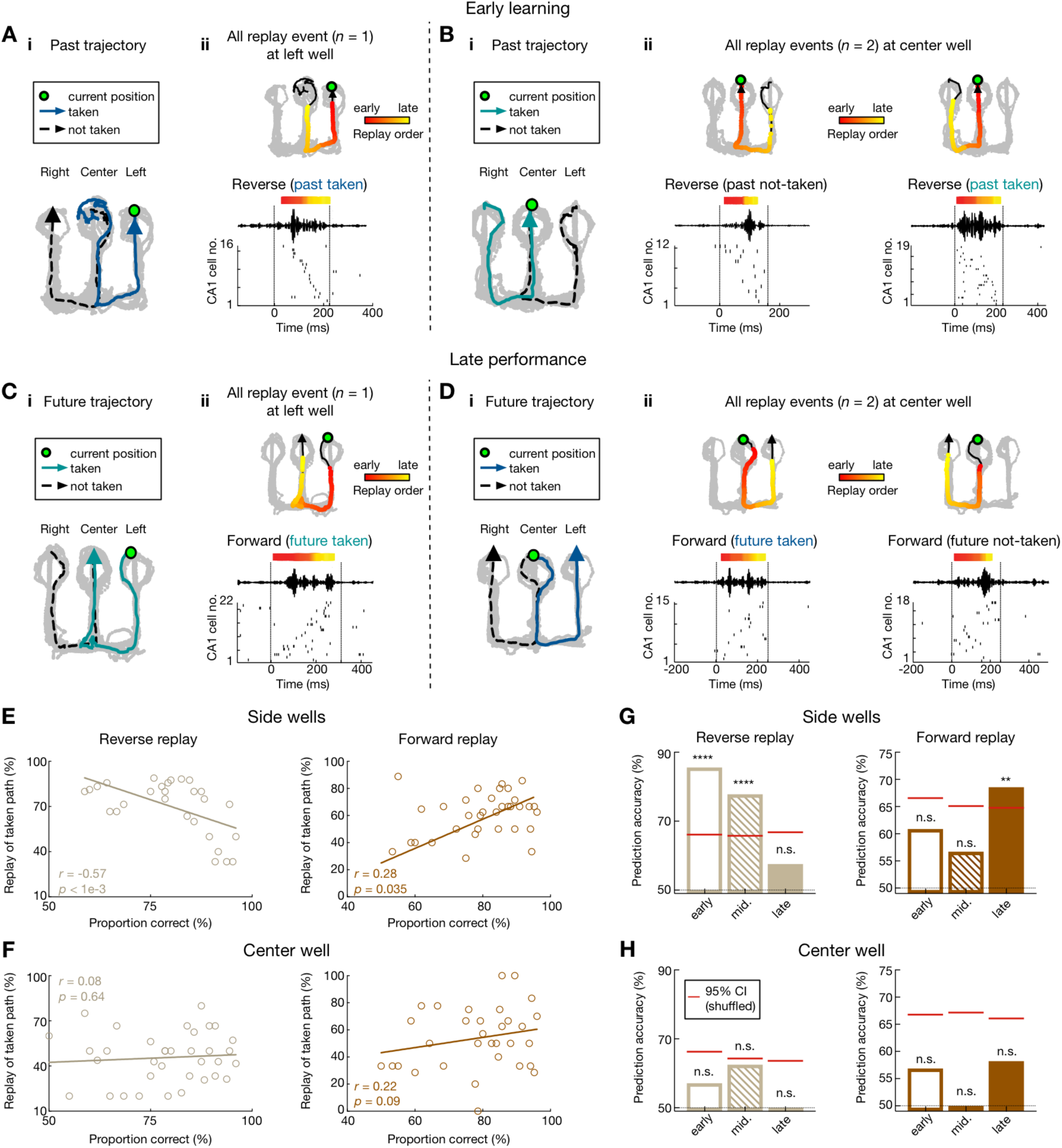
Contrasting evolution of reverse and forward replay during the course of learning. **(A** and **B)** Example reverse replay events during early learning at **(F)** the side well and **(G)** the center well. Data are presented as in **Figure 3**. **(C** and **D)** Example forward replay events during late performance at **(H)** the side well and **(I)** the center well. See also **Figure S4** for additional replay examples. **(E** and **F)** Relationship between task performance and fraction of reverse replay of past taken paths (*left*), or forward replay of future taken paths (*right*) at **(E)** the side wells and **(F)** the center well. Each dot represents one session from a subject. Note the significant negative correlation for reverse replay (*r* = −0.57, *p* < 0.0001) and positive correlation for forward replay (*r* = 0.28, *p* = 0.035; permutation tests) at the side wells, but not the center well (*r* = 0.08, *p* = 0.64 for reverse replay; *r* = 0.22, *p* = 0.09 for forward replay). **(G)** At the side wells, reverse replay events predict taken past paths during early learning sessions (*left*), while forward replay events predict future taken paths during late performance sessions (*right*). Cross-validated SVM decoders were trained during early, middle and late sessions (sessions 1-3, 4-5, and 6-8, respectively) based on the number of either forward or reverse replay events of 4 possible trajectories at the wells during correct trials. Only trials with at least one replay event of the given type (forward or reverse) were used (*n* = 79, 85, and 85 trials for reverse, and *n* = 87, 95, and 104 trials for forward during early, middle and late sessions, respectively). Significant prediction power was only observed during early and middle sessions for reverse replay (*left*; *p* < 0.0001****, 0.0001**** and 0.43 for early, middle and late sessions, respectively), and late sessions for forward replay (*right*; *p* = 0.23, 0.35, and 0.003** for early, middle and late sessions, respectively). **(H)** Taken paths cannot be predicted from the patterns of replay events at the center well (*n* = 89, 91, and 118 trials, and *p* = 0.36, 0.08, and 0.68 for reverse replay; *n* = 72, 71, and 74 trials, and *p* = 0.38, 0.56, and 0.15 for forward replay during early, middle and late sessions, respectively). Red horizontal lines on columns represent chance levels calculated by permutation tests.

Replay events therefore shifted their content over the course of learning at the side wells, which was apparent both in correlations of replay content with behavioral performance, as well as the ability of replay to predict behaviorally actualized past and future paths (**Figure 5G**; overall, forward replay events of future taken vs. not-taken paths: 169/387 vs. 103/387, 43.7% vs. 26.7%, *p* < 1e-4; reverse replay events of past taken vs. not-taken paths: 200/376 vs. 67/376, 53.2% vs. 17.7%, *p* < 1e-4, z-test for proportions). Reverse replay content was able to accurately predict the actual past path of the animal during early stages of learning, but not later performance (**Figure 5G**, *Left*; cross-validated SVM decoders were trained on the number of reverse replay events to predict taken paths, see **STAR Methods**). Forward replay of taken future paths showed significant positive correlation with behavioral performance (**Figure 5E**, *Right*), with accurate prediction of actual future path emerging only after learning during later performance sessions (**Figure 5G**, *Right*). This change in replay content from reverse replay of past taken path to forward replay of future upcoming path of the animal thus underscores a learning shift from retrospective evaluation of the completed outbound trajectory terminating in reward, to prospective planning of the future inbound trajectory that underlies execution of the reference memory rule.

In marked contrast, replay events at the center well, marking the transition point of the history-dependent spatial working memory behavioral sequence, did not show a preferential bias towards actual taken paths, either for reverse replay of past paths (**Figure 5F**, *Left*; overall, reverse replay events of past taken vs. not-taken paths: 146/439 vs. 177/439, 33.6% vs. 40.3%), or for forward replay of future paths (**Figure 5F**, *Right*; overall, forward replay events of future taken vs. not-taken paths: 111/293 vs. 91/293, 37.9% vs. 31.6%; *p* = 0.55, z-test for proportions). Reverse replay content did not show correlation with performance and could not predict the actual taken past path that terminated at the center well for any learning stage (**Figure 5H**, *Left*). Similarly, forward replay content was unable to predict the actual taken future path that originated at the center well (**Figure 5H**, *Right*). Thus, rather than a deterministic role of hippocampal replay in representing past and future paths comprising a spatial working-memory sequence, this suggests a persistent evaluative role of replay throughout the course of learning and performance. Hippocampal replay can therefore underlie a cognitive exploration of possible past and future paths (Gupta et al., 2010; Redish, 2016; Singer et al., 2013; Stella et al., 2019) for this history-dependent spatial working-memory rule, and we hypothesized that selection of behaviorally relevant replayed trajectories for this deliberative working memory process (i.e. choosing correct future trajectory based on executed past trajectory) occurs in networks outside the hippocampus, with the prefrontal cortex a likely candidate (Eichenbaum, 2017; Shin and Jadhav, 2016; Spellman et al., 2015; Tang and Jadhav, 2018; Yu and Frank, 2015).

### Coordinated hippocampal-prefrontal replay selects past-future trajectory sequences

We therefore examined the relationship of coherent hippocampal-prefrontal (CA1-PFC) replay of spatial paths to ongoing behavioral trajectories (Tang et al., 2017). Similar to CA1 (**Figure 1**), PFC neurons exhibited spatially and directionally selective firing, with PFC ensembles forming unique spatial representations of different trajectories on the maze for all sessions (**Figure 6**). We and others have shown that although PFC neurons have significantly lower spatial specificity and multi-peaked fields as compared with CA1 neurons (Jadhav et al., 2016; Tang et al., 2017; Yu et al., 2018; Zielinski et al., 2019), PFC ensembles can still represent spatial location with high accuracy (Mashhoori et al., 2018; Zielinski et al., 2019). Just as in CA1, spatial- and directional-selective firing in PFC was seen across all 8 sessions starting with the first session on the track (**Figures 6A** and **6B**). PFC ensembles thus represent unique locations along different trajectories on the maze for all sessions – their activity accurately predicted animal position during spatial exploration using a Bayesian decoder for individual sessions (**Figures 6B-6E**) (Tang et al., 2017; Zielinski et al., 2019), and ensemble activity exhibited significant directionality for all 8 sessions (**Figures 6A, 6B** and **6F**).

**Figure 6.**
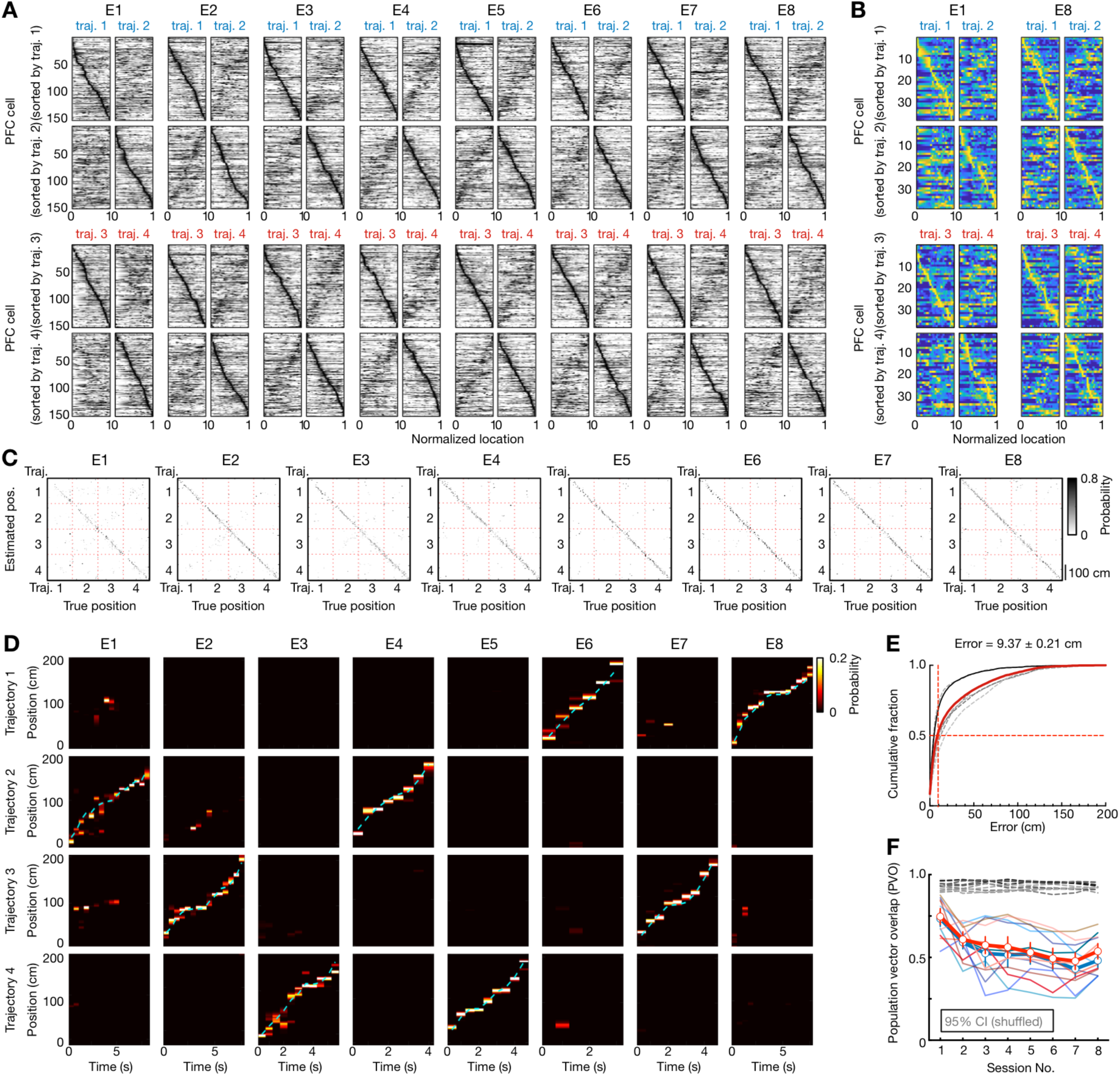
Spatial coding and distinct representations of behavioral trajectories by prefrontal cells. **(A)** Spatial maps of all PFC cells (*n* = 154) recorded from 6 rats continuously across 8 learning sessions (E1-8). Data are presented as Figure 1C. Note that patterns between pairs of trajectories reflect path equivalence properties of PFC neurons (Yu et al., 2018). **(B)** Spatial maps of all PFC cells (*n* = 38) recorded from an example animal in the first and last sessions. **(C-E)** Position reconstruction based on PFC ensemble spiking from the example animal in **(B)** during active running. **(C)** Confusion matrices (estimated vs. true position). **(D)** Estimated position probabilities. Cyan line: actual animal trajectory. Data are presented as Figures 1D and 1E. **(E)** Cumulative PFC decoding errors across all animals (*n* = 6 rats). Dashed lines: individual animals; Red line: all animals; Solid black line: the example animal shown in **(B-D)**. Median error of all sessions noted on top as (median ± SEM). **(F)** Population vector overlap (PVO) of PFC population activity in two running directions across 8 learning sessions. Data are presented as in **Figure 1H.** Note that the spatial maps of PFC population in two running directions became less similar across sessions, but distinct templates for each direction were apparent as early as the first session (*p’s* < 0.0001 compared to the shuffled data for individual rats, permutation tests).

We have previously reported coordinated hippocampal-prefrontal replay during SWRs (Jadhav et al., 2016; Tang et al., 2017), and therefore examined the relationship between coherent hippocampal-prefrontal replay and behavioral choices. Here, coherent CA1-PFC reactivation is defined as a CA1 replay event where the same trajectory is also significantly reactivated by CA1-PFC ensembles, detected as ‘reactivation strength’ using a template matching method, similar to previous reports (**Figure 7**; **Figures S7** and **S8**; see **STAR Methods**; a comparison of the template matching method with Bayesian decoding and line-fitting methods is detailed in **Figure S8**, using both model simulations and experimental data) (Girardeau et al., 2017; Lansink et al., 2009; Peyrache et al., 2009; Tang et al., 2017). Using template spatial maps for CA1 and PFC neurons and candidate coherent replay events (≥ 5 PFC and CA1 place cells active; **Figures 7A-7D**), we calculated the ensemble CA1-PFC correlation coefficient during running behavior (C_RUN_) and SWR events (C_SWR_) for each of the four possible trajectories. The reactivation strength of each of these events was measured as the correlation between the population matrices, C_RUN_ and C_SWR_. Illustrative coherent CA1-PFC reactivation events, with both forward and reverse CA1 replay, are shown in **Figures 7A-7D**, with the linearized spatial maps of each cell that fired during the SWR replay event (**Figures 7A_i_-7D_i_**), the activity for a single running trial and activity during the replay event (**Figures 7A_ii_-7D_ii_**), the corresponding replayed trajectory in CA1 (**Figures 7A_iii_-7D_iii_**), and the reactivation strength of the coherent CA1-PFC replay trajectory (**Figures 7A_iv_-7D_iv_**). Crucially, this enabled us to compare the strength of coherent CA1-PFC replay when CA1 replayed either the behaviorally taken path, or the alternative not-taken path as a paired comparison during each event. We found that coherent CA1-PFC reactivation was significantly stronger when CA1 replayed the actual taken paths as compared with the alternative trajectory, but not when CA1 replayed the not-taken paths (**Figure 7E**; effect seen at both center and side wells). Stronger coherent CA1-PFC replay was true for both forward CA1 replay of future taken paths, as well as reverse CA1 replay of past taken paths (**Figure 7F**). The degree of coherent CA1-PFC replay (quantified as the fraction of coherent CA1-PFC replay events) was also higher for taken as compared with not-taken CA1 replay (**Figure 7G**).

**Figure 7.**
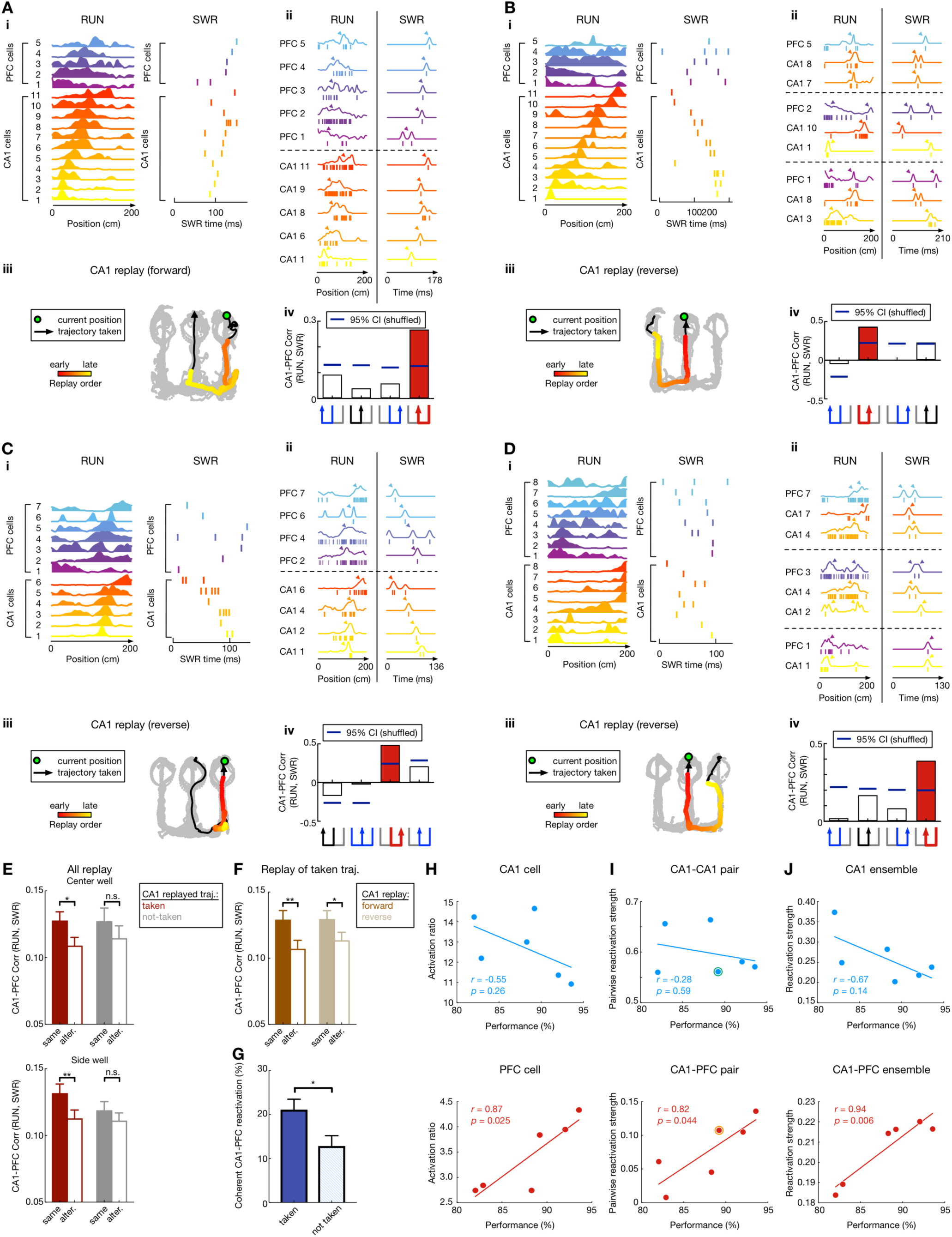
Coherent hippocampal-prefrontal (CA1-PFC) reactivation of past and future trajectories. **(A-D)** Four examples of coherent CA1-PFC reactivation of future and past taken paths. **(i)** Linearized CA1-PFC spatial-map template with cell IDs on y-axis (*left*), and the corresponding raster plot during the SWR (*right*). **(ii)** Detailed view of CA1-PFC coordination during RUN and the SWR using example cells. *Left* (RUN): the spiking pattern from a single running trial (ticks) with linearized spatial maps obtained by averaging over all trials (overlaid lines). *Right* (SWR): spikes (ticks) during the SWR, and the response curves (overlaid lines) created by smoothing observed spikes with a Gaussian kernel. Arrowheads indicate the peak locations. **(ii)** CA1 trajectory replay, with data presented as in Figure 5A. Green circle: animal’s current position; Black arrowhead line: trajectory taken; Colored dots: decoded CA1-replay path. **(iv)** Reactivation strength, measured as the correlation coefficient of CA1-PFC activity during RUN vs. SWR using template matching, for four possible trajectories (trajectory schematic on the bottom; red for CA1-replay path, black for alternative path, blue for the other paths). Blue horizontal lines on columns represent 95% confident intervals computed from shuffled data (SWR spike time shuffle). Selected PFC cells with highest contribution to the reactivation strength are shown in **(iii)** for ease of presentation, illustrating the synchronized firing pattern for CA1 and PFC cells during RUN, which is reactivated during the SWR event. **(E)** Paired comparison of CA1-PFC reactivation strength for CA1-replay of taken vs. alternative path during all CA1 replay events at center (*Top*) and side (*Bottom*) wells. Note that CA1-PFC reactivation strength is significantly higher for the CA1-replayed path compared to its alternative, only when this path was the behaviorally taken path at both center (*p* = 0.032*) and side (*p* = 0.0075**) wells, but not when the CA1-replayed path was the “not-taken” path (*p* = 0.35 and 0.16 for side and center wells, respectively; rank-sum paired tests). **(F)** Stronger CA1-PFC reactivation of taken path compared to not-taken path for both forward and reverse CA1 replay (*p* = 0.0058** and 0.10 for forward replay of taken and not-taken, respectively; *p* = 0.039* and 0.65 for reverse replay of taken and not-taken, respectively; rank-sum paired tests). **(G)** The proportion of coherent CA1-PFC reactivation events is significantly higher during CA1 replay of taken vs. not-taken trajectories (*p* = 0.02*, session-by-session rank-sum paired test). **(H-J)** Relationship between animals’ maximal performance level on the outbound, spatial working memory component and **(H)** activation ratio of CA1 (*Top*) and PFC (*Bottom*) cells during SWR synchronous events, **(I)** reactivation strength of CA1-CA1 (*Top*) and CA1-PFC (*Bottom*) cell pairs (open circle indicates the example animal shown in **Figure S7G**), and **(J)** reactivation strength of CA1 (*Top*) and CA1-PFC (*Bottom*) populations (see **STAR Methods**). Each dot represents an animal. Behavioral performance showed significant correlation with PFC activation ratio, CA1-PFC pairwise, and ensemble reactivation strength, but not for CA1-only parameters. See also **Figures S7** and **S8**.

Coherent CA1-PFC replay thus mediated selection, through stronger reactivation, for actual (i.e., behaviorally instantiated) past and future paths during reverse and forward CA1 replay, respectively (**Figures 4E-4G**). This was not a result of any difference in CA1 replay quality for taken vs. not-taken paths (**Figure S7A**). Interestingly, we also observed that stronger coherent CA1-PFC replay for future taken paths vs. not-taken paths emerged only during late performance stages, when animals performed well above chance-levels (**Figure S7C**). This suggests emergence of prospective planning of correct upcoming choices through coherent CA1-PFC replay as animals learn the task.

Finally, we asked if coherent CA1-PFC replay was related to behavioral performance of animals, and indeed found a significant correlation between higher CA1-PFC reactivation and working-memory performance (**Figures 7H-7J**; examples in **Figures S7E-S7G**). The degree of engagement of PFC activity (and not CA1 activity) during SWR replay events, the strength of pairwise CA1-PFC reactivation (and not within-CA1 pairwise reactivation), and the magnitude of CA1-PFC ensemble reactivation (and not CA1 ensemble reactivation) corresponded to maximal outbound behavioral performance across the six animals, suggesting a relationship between CA1-PFC replay and memory performance (**Figures 7H-7J**).

## DISCUSSION

Our results provide novel insights into the role of replay in spatial memory, and suggest a mechanism of coordinated hippocampal-prefrontal replay underlying retrospective evaluation for learning in novel environments, and prospective planning for decision making after task acquisition to support memory performance. Continuous tracking of replay over the course of spatial choice learning revealed that reverse replay mediates retrospective evaluation of possible past paths leading to goals, and forward replay mediates prospective planning of available future choices toward goals. Crucially, we found dynamic changes in functional roles of replay depending on the learning stage, as well as a novel role in mediating spatial working memory tasks that requires coherent hippocampal-prefrontal reactivation. The W-track alternation task comprises interleaved components of history-independent (return-to-center trajectory) and history-dependent (side-to-center followed by center-to-opposite-side trajectory sequence) spatial rules. Our results suggest a model (**Figure 8**) with differing roles of replay in (i) history-dependent spatial working-memory tasks which necessarily need coherent hippocampal-prefrontal replay for recall of the actual past experience to guide selection of the future choice; and (ii) history-independent, deterministic spatial rules where the outcome is pre-determined based only on current location, similar to previous studies with pre-determined paths between reward wells (Ambrose et al., 2016; Diba and Buzsáki, 2007; Foster and Wilson, 2006; Ólafsdóttir et al., 2016, 2017; Wu et al., 2017).

**Figure 8.**
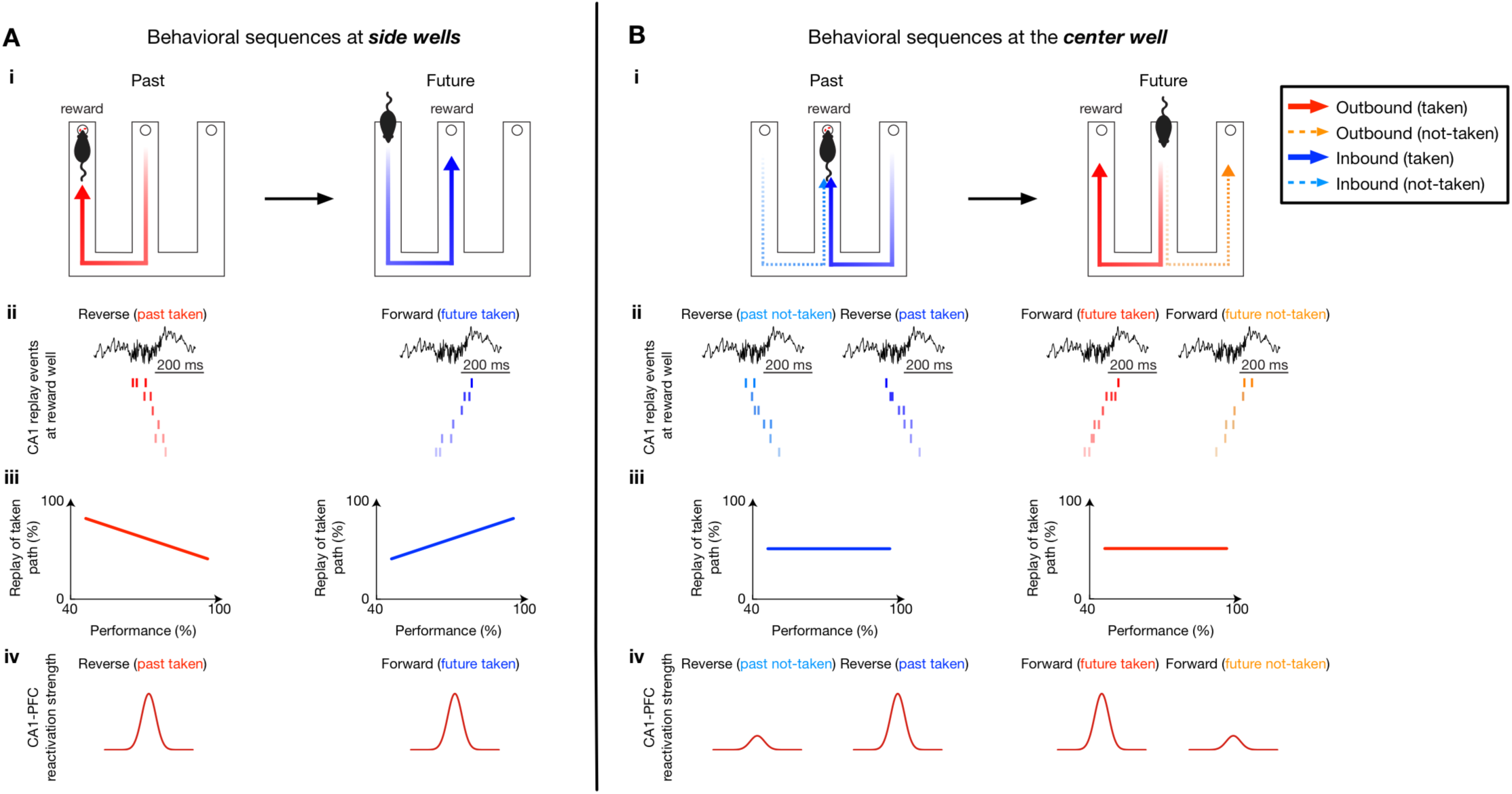
Schematic: hippocampal-prefrontal replay mediates retrospection and prospection for learning and planning in spatial memory-guided tasks. **(A)** The role of hippocampal-prefrontal replay in learning history-independent, deterministic spatial rules. **(A_i_)** At the side-wells, past paths are outbound components (center-to-side), and future paths are inbound components (side-to-center). **(A_ii_-A_iii_)** Hippocampal replay showed a shift from reverse-replay-based prediction of taken past path during early learning, to forward-replay-based prediction of taken future path during late performance, along with **(A_iv_)** stronger coherent CA1-PFC reactivation of taken paths. **(B)** The role of hippocampal-prefrontal replay in learning history-dependent spatial working-memory rules. **(B_i_)** At the center well, past paths are inbound components (side-to-center), and future paths are outbound components (center-to-side), with a past-future trajectory sequence spanning the center well transition sub-serving a replay-dependent spatial working memory task (Jadhav et al., 2012). **(B_ii_)** Hippocampal replay persistently reverse-replayed both past choices and forward-replayed both future choices at the center well **(B_iii_)** throughout the course of learning and performance. **(B_iv_)** For each CA1 replay event, coherent CA1-PFC replay discriminates the behaviorally taken past path and the chosen future path from the alternatives, and can mediate planning of correct future trajectory based on the completed past trajectory.

At the side wells, replay showed a shift from reverse-replay-based prediction of past path during early learning, to forward-replay-based prediction of future path during late performance. Reverse and forward hippocampal replay representations of past and future paths are in agreement with observations for deterministic spatial trajectories on linear tracks between reward wells (Ambrose et al., 2016; Diba and Buzsáki, 2007; Foster and Wilson, 2006), and our results establish a learning gradient for the role of reverse and forward replay (**Figure 8A**). Specifically, for forward replay, our findings indicate that with repeated experience of the same trajectory spanning reward wells in the inbound return-to-center reference memory component, prediction of future path leading to reward emerges over learning. This forward replay prediction is in agreement with previous observations in spatial reference memory tasks (Pfeiffer and Foster, 2013; Wu et al., 2017; Xu et al., 2019), and emergence of this prediction with learning has been hypothesized (Pfeiffer, 2018). Interestingly, disrupting hippocampal replay does not impair learning for the inbound component in the W-track task (Jadhav et al., 2012), suggesting that despite the prediction ability of forward replay, other mechanisms can potentially support (or compensate) learning of deterministic paths leading to goals.

Reverse replay at side wells mediated retrospection of the past outbound (center-to-side) paths leading to reward. This reverse replay prediction occurs at the completion of the spatial working memory trajectory, and prediction of the past path leading to reward aligns with a role in spatial working memory updates reported in a radial-arm maze task (Xu et al., 2019). Intriguingly, we found that this bias toward taken past path is present only during early learning. This indicates that reverse replay of goal-directed past paths can play an important role in temporal credit assignment during early learning in a novel environment (Foster and Knierim, 2012; Haga and Fukai, 2018; Mattar and Daw, 2018; Pfeiffer, 2018), and the loss of bias for taken past path over learning supports the hypotheses that this credit assignment function is no longer required after acquisition of the task (Foster and Knierim, 2012). Notably, despite the findings of reverse-past-replay and forward-future-replay, a simple model of latent excitability (Atherton et al., 2015; Battaglia et al., 2011; Csicsvari et al., 2007) is insufficient to explain reverse and forward directions of replay, since they occur in an inter-mixed order during immobility and not just at beginning and end of immobility periods, which has also been reported in a previous study (Ambrose et al., 2016). Finally, stronger coherent CA1-PFC reactivation was observed for taken past and future paths at the side wells, suggesting behavioral mediation by prefrontal reactivation.

In contrast, surprisingly, hippocampal replay exclusively underlies a lower-level cognitive search role, and not a predictive element, for working memory rules (**Figure 8B**). At the center well, past paths are inbound components (side-to-center), and future paths are outbound components (center-to-side), with a past-future trajectory sequence spanning the center well transition sub-serving a replay-dependent spatial working memory task (Jadhav et al., 2012). We found that hippocampal replay persistently reverse-replayed both possible past choices and forward-replayed both available future choices throughout the course of learning and performance. The lack of bias toward taken paths is not simply an artifact of possible issues in replay detection, as demonstrated by the learning gradient of trajectory prediction for side wells, providing an internal control. This is indicative of a priming process for retrospection and prospection (Buzsáki, 2015), with hippocampal replay underlying a cognitive exploration of possible paths (Gupta et al., 2010; Pfeiffer, 2018; Redish, 2016; Singer et al., 2013; Stella et al., 2019) that can be utilized by other networks for deliberation and prospective planning (**Figure 8B**). For individual hippocampal replay events, coherent hippocampal-prefrontal replay can discriminate the behaviorally taken past path and the chosen future path from the alternatives, suggesting that it supports recall of the actual past experience and selection of the future choice based on the hippocampal evaluative process. Coherent CA1-PFC replay can thus mediate planning of correct future trajectory based on the completed past trajectory. This PFC read-out interpretation is supported by a bias toward CA1-leading-PFC directionality during replay (Jadhav et al., 2016; Rothschild et al., 2016), which can potentially be tested using selective causal perturbation of coherent PFC replay (Shin and Jadhav, 2016; Tang and Jadhav, 2018; Zielinski et al., 2017).

Our results thus suggest a novel role of replay in acquisition and performance of history-dependent spatial working memory rules. Hippocampal replay of possible choices underlies an evaluative process for retrospection and prospection respectively, mediating a cognitive exploration of possible paths, as previously hypothesized (Colgin, 2016; Foster, 2017; Gupta et al., 2010; Jadhav et al., 2012; Redish, 2016; Singer et al., 2013; Stella et al., 2019), that can be utilized by other networks for reinforcement learning and prospective planning. Coherent CA1-PFC replay distinguishes behaviorally taken past and future paths during the spatial working memory sequence, supporting a key role of PFC in trajectory choice selection for spatial working memory (Jadhav et al., 2012; Spellman et al., 2015; Tang and Jadhav, 2018; Yu and Frank, 2015). We hypothesize that coherent replay of the future planned path can influence the behavioral time-scale, choice-selective activity (Fujisawa et al., 2008; Guise and Shapiro, 2017; Ito et al., 2015; Zielinski et al., 2019) that is seen in these networks as animals execute the chosen future action and run toward reward. It is important to point out that our experimental design enabled a rapid learning time-scale, and it is possible that this replay pattern is not seen in repeatedly trained tasks, when other habitual systems can contribute to learning and performance (Kim and Frank, 2009; Packard and McGaugh, 1996).

This model of hippocampal cognitive exploration for selection and filtering by prefrontal (and possibly other) networks has implications for neural mechanisms of model-based learning and planning, possibly for spatial as well as non-spatial memories (Daw et al., 2005; Doll et al., 2012; Miller et al., 2017; Pezzulo et al., 2014; Redish, 2016; Smittenaar et al., 2013; Vikbladh et al., 2019). In the W-track task, multiple paths leading to and from the center well underlie a non-Markovian structure (Mattar and Daw, 2018), and the spatial working-memory component of the task requires animals to integrate across space and time to learn sequences of past and future trajectory choices that lead to reward. Hippocampal replay events consistently provide a cognitive exploration of past and future choices via reverse and forward replay, a process that can be influenced or read-out by inputs and outputs from other regions (Gomperts et al., 2015; Ji and Wilson, 2007; Pfeiffer, 2018; Tang and Jadhav, 2018; Yamamoto and Tonegawa, 2017). We suggest that replay in the awake state represents an internal cognitive state that engages a broad, multi-region network, similar to a default network mode (Buckner, 2010; Logothetis et al., 2012), to support ongoing learning, prospection, and imagination.

## ACKNOWLEDGMENTS

This work was supported by NIH Grant R01 MH112661, a Sloan Research Fellowship in Neuroscience (Alfred P. Sloan Foundation), and Whitehall Foundation award to S.P.J.

## AUTHOR CONTRIBUTIONS

J.D.S, W.T. and S.P.J. conceived and designed the study, collected and analyzed the data, and wrote the manuscript. J.D.S. and W.T. contributed equally to this study.

## STAR ★ METHODS

### KEY RESOURCES TABLE

| REAGENT or RESOURCE | SOURCE | IDENTIFIER |
| --- | --- | --- |
| Chemicals |  |  |
| Cresyl Violet | Acros Organics | Cat#: AC229630050 |
| Formaldehyde | Fisher | Cat#: 50-00-0,67561,7732-18-5 |
| Isoflurane | Patterson Veterinary | Cat#: 07-806-3204 |
| Ketamine | Patterson Veterinary | Cat#: 07-803-6637 |
| Xylazine | Patterson Veterinary | Cat#: 07-808-1947 |
| Atropine | Patterson Veterinary | Cat#: 07-869-6061 |
| Bupivacaine | Patterson Veterinary | Cat#: 07-890-4881 |
| Beuthanasia-D | Patterson Veterinary | Cat#: 07-807-3963 |
| Sucrose | Sigma-Aldrich | Cat#: S8501-5KG |
| Experimental Models: Organisms/Strains |  |  |
| Rat: Long Evans | Charles River | Cat#: Crl:LE 006;<br>RRID:<br>RGD 2308852 |
| Software and Algorithms |  |  |
| MATLAB 2017a | Mathworks, MA | RRID: SCR_001622 |
| Trodes | SpikeGadgets | http://www.spikegadgets.com |
| Matclust | Mattias P. Karlsson | https://www.mathworks.com/matlabcentral/fileexchange/39663-matclust, V1.7 |
| Libsvm | Chang and Lin 2011 | https://www.csie.ntu.edu.tw/~cjlin/libsvm/, V3.12 |
| Prism 8 | GraphPad Software | RRID: SCR_002798 |
| Other |  |  |
| 128 Channel electrophysiology data acquisition system | SpikeGadgets | http://www.spikegadgets.com |
| 12.7 μm NiCr tetrode wire | Sandvik | Cat#: PX000004 |

### CONTACT FOR REAGENT AND RESOURCE SHARING

Further information and requests for resources should be directed to and will be fulfilled by the Lead Contact, Dr. Shantanu P. Jadhav.

### EXPERIMENTAL MODEL AND SUBJECT DETAILS

All procedures were approved by the Institutional Animal Care and Use Committee at the Brandeis University and conformed to US National Institutes of Health guidelines. Six adult male Long-Evans rats (450-550 g, 4-6 months) were used in this study. Animals were individually housed and kept on a 12-hr regular light/dark cycle.

### METHOD DETAILS

#### Experimental design

Prior to training, animals were first habituated to daily handling over several weeks. After habituation, animals were food deprived to 85-90% of their *ad libitum* weight and pre-trained to seek liquid food rewards (sweetened evaporated milk) at each end of an elevated linear track by running back and forth as described previously (Jadhav et al., 2012). After pre-training on linear-track behaviors and habituation to an elevated, opaque sleep box, animals were surgically implanted with a multi-tetrode drive (see *Surgical implantation and electrophysiology*). Following recovery from surgical implantation (∼7-8 d), animals were food-deprived and again pre-trained on a linear track for at least two days before the W-maze sessions started. During the experimental day, animals were introduced to the novel W-maze for the first time during the recording sessions, and learned the task rules over 8 behavioral sessions (see below, *The W-maze spatial memory task*). Following the conclusion of the experiments, micro-lesions were made through each electrode tip to mark recording locations (Jadhav et al., 2012; Jadhav et al., 2016). After 12-24 hours, animals were euthanized (Beuthanasia) and intracardially perfused with 4% formaldehyde using approved procedures. Brains were fixed for 24 hours, cryoprotected (30% sucrose in 4% formaldehyde), and stored at 4 °C. The recording sites were determined from *post hoc* Nissl-stained coronal brain sections based on *The Rat Brain in Stereotaxic Coordinates* (Paxinos and Watson, 2004) (**Figure S1**).

#### The W-maze spatial memory task

Animals learned a novel W-maze continuous spatial alternation task within a single day. During this experimental day, all animals ran eight 15-20 min sessions on a W-maze interleaved with 20-30 min rest sessions in a sleep box (W-maze sessions: 17.9 ± 1.0 mins per session, 8 sessions per rat; rest sessions: 23.0 ± 4.9 mins per session, 9 sessions per rat; total recording duration: 6.04 ± 0.37 hrs per rat; mean ± SD). The W-maze was novel in the first behavioral session (sleep box was opaque, and the animal had no visibility of the W-maze until it was introduced in the first run session), and had dimensions of ∼ 80 × 80 cm with ∼7-cm-wide track sections. Three reward food wells (i.e., right, center and left wells) were located at the end of three arms of the W-maze (**Figure 1**). Calibrated evaporated milk reward was automatically delivered in the reward wells triggered by crossing of an infrared beam by the animal’s nose. Rewards were delivered according to the following rules (**Figure 1A**): returning to the center well after visits to either side well (inbound trajectories), and choosing the opposite side well from the previously visited side well when starting from the center well (outbound trajectories). Incorrect alternations (visiting the same side well in consecutive outbound components – outbound error), or incorrect side-to-side well visits (without visiting the center arm – inbound error) were not rewarded. Repeated visits to the same well were also not rewarded (i.e., turn-around error). Therefore, animals performed four types of trajectories during correct behavioral sequences (**Figure 1A**): center-to-right (C-to-R), right-to-center (R-to-C), center-to-left (C-to-L) and left-to-center (L-to-C). Among these trajectory types, C-to-R and C-to-L are outbound trajectories, while R-to-C and L-to-C are inbound trajectories. When animals paused at one reward well during correct trials, two of these four trajectory types represented the immediate past and future paths taken, and the other two represented the alternative not-taken paths (**Figure 1B**). For visualization purposes, the alternative, not-taken trajectories corresponding to a taken behavioral sequence were selected from the adjacent trials (e.g., **Figures 3D** and **3H**). Only behaviorally correct trials were included for replay and reactivation analyses, unless otherwise specified (see below, *Neural Analyses*). At the end of each W-maze session, animals were transferred to a black opaque box for rest (∼ 30 × 30 cm with a 50-cm high wall). The raw performance of the task was calculated as proportion correct (Singer et al., 2013) (**Figure 4A**) and the learning curves were estimated using a state-space model (Jadhav et al., 2012; Smith et al., 2004) (**Figure S2**). Each animal’s performance level (**Figures 7H-7J**) was measured as the highest performance reached on the outbound learning curve (**Figure S2**). All 6 animals performed > 80% correct in the W-maze task toward the end of learning (maximal proportion correct of outbound for individual animals: 91.2 ± 4.1%; mean ± SD).

#### Surgical implantation and electrophysiology

All 6 animals performed > 80% correct in the W-maze task toward the end of learning (maximal proportion correct of outbound for individual animals: 91.2 ± 4.1%; mean ± SD). Surgical implantation procedures were as previously described (Jadhav et al., 2012; Jadhav et al., 2016; Tang et al., 2017). Animals were implanted with a microdrive array containing 30-32 independently moveable tetrodes targeting right dorsal hippocampal region CA1 (−3.6 mm AP and 2.2 mm ML) and right PFC (+3.0 mm AP and 0.7 mm ML). On the days following surgery, hippocampal tetrodes were gradually advanced to the desired depths with characteristic EEG patterns (sharp wave polarity, theta modulation) and neural firing patterns as previously described (Jadhav et al., 2012; Jadhav et al., 2016). One tetrode in corpus callosum served as hippocampal reference, and another tetrode in overlying cortical regions with no spiking signal served as prefrontal reference. A ground (GND) screw installed in skull overlying cerebellum also served as a reference. All spiking activity and ripple-filtered LFPs (150-250 Hz; see below) were recorded relative to the local reference tetrode. Electrodes were not moved at least 4 hours before and during the recording day.

Data were collected using a SpikeGadgets data acquisition system (SpikeGadgets LLC) (Tang et al., 2017). Spike data were sampled at 30 kHz and bandpass filtered between 600 Hz and 6 kHz. LFPs were sampled at 1.5 kHz and bandpass filtered between 0.5 Hz and 400 Hz. The animal’s position and running speed were recorded with an overhead color CCD camera (30 fps) and tracked by color LEDs affixed to the headstage.

Spiking activity was continuously monitored during the experimental day for ∼6-7 hrs. Single units were identified by manual clustering based on peak and trough amplitude, principal components, and spike width using custom software (MatClust, M. P. Karlsson) as previously described (Jadhav et al., 2016; Tang et al., 2017). Only well isolated neurons with stable spiking waveforms were included (**Figure S1**). Cluster quality was assessed using isolation distance (Schmitzer-Torbert et al., 2005), cluster center-of-mass shift (Mallory et al., 2018), and spike-waveform correlation (Li et al., 2017) (**Figure S1**). Cluster center-of-mass shift between two different sessions was calculated as the Mahalanobis distance between the cluster centroids of the same single unit from these sessions. The spike-waveform correlation was quantified as the correlation coefficient between averaged spike waveforms of the same single unit from two consecutive sessions, and the resulting correlation was Fisher-transformed to make it normally distributed (Li et al., 2017).

#### Unit inclusion

Units included in analyses fired at least 100 spikes in each session. Putative interneurons were identified and excluded based on spike width and firing rate criterion as previously described (Jadhav et al., 2016; Tang et al., 2017). Peak rate for each unit was defined as the maximum rate across all spatial bins in the linearized spatial map (see *Spatial maps*). A peak rate ≥ 3 Hz was required for a cell to be considered as a place cell. Only cells recorded continuously across all 8 behavioral sessions with stable spiking waveforms were analyzed (**Figure S1**).

#### Behavioral state definition

Movement or exploratory states were defined as periods with running speed > 5 cm/s, whereas immobility was defined as periods with speed ≤ 4 cm/s. The animal’s arrival and departure at a reward well was detected by an infrared beam triggered at the well. The well entry was further refined as the first time point when the speed fell below 4 cm/s before the arrival trigger, whereas the well exit was defined as the first time point when the speed rose above 4 cm/s after the departure trigger (**Figures 3A** and **3D**). The animal’s time spent at a reward well (i.e., immobility period at well) was defined as the period between the well entry and exit.

### QUANTIFICATION AND STATISTICAL ANALYSIS

#### Sharp-wave ripple detection

SWRs were detected as described previously during immobility periods (≤ 4 cm/s) (Jadhav et al., 2016; Karlsson and Frank, 2009; Tang et al., 2017). In brief, LFPs from CA1 tetrodes were filtered into the ripple band (150-250 Hz), and the envelope of the ripple-filtered LFPs was determined using a Hilbert transform. SWRs were initially detected as contiguous periods when the envelope stayed above 3 SD of the mean on at least one tetrode, and further refined as times around the initially detected events during which the envelope exceeded the mean. For replay and reactivation analysis (see below, *Replay decoding* and *CA1-PFC reactivation analysis*), only SWRs with a duration ≥ 50 ms were included, similar to previous studies (Pfeiffer and Foster, 2013; Wu et al., 2017).

#### Spatial maps

Spatial maps were calculated only during movement periods (> 5 cm/s; all SWR times excluded) at positions with sufficient occupancy (> 20 ms). Two-dimensional occupancy-normalized spatial rate maps were calculated as previously described (Jadhav et al., 2012; Jadhav et al., 2016; Tang et al., 2017). To construct the spatial-map templates of different trajectory types on a W-maze (**Figure 1C**), we calculated the linearized activity of each cell as previously described (Jadhav et al., 2012; Jadhav et al., 2016; Karlsson and Frank, 2009; Singer et al., 2013). The rat’s linear position was estimated by projecting its actual 2D position onto pre-defined idealized paths along the track, and was further classified as belonging to one of the four trajectory types. The linearized spatial maps were then calculated using spike counts and occupancies calculated in 2-cm bins of the linearized positions and smoothened with a Gaussian curve (4-cm SD) as previously described (Jadhav et al., 2012; Tang et al., 2017). To cross-validate the linearized positions, an alternative linearization method was also used based on nearest-neighbor Delaunay triangulation (Ferbinteanu et al., 2011). Completed trials that were detected based on both methods, i.e., linearized trajectories starting from and ending at reward wells, were used for replay and reactivation analyses. To quantify spatial coverage of place-cell populations, a spatial bin was considered as represented if at least one cell from the population had an occupancy-normalized rate ≥ 3 Hz within the bin (Kay et al., 2016; Zielinski et al., 2019). The spatial coverage of the population was then expressed as the percentage of the spatial bins covered. Across the populations of recorded place cells, we found place fields at all positions along each trajectory type (spatial coverage per subject over sessions, shown as mean ± SD: 99.9 ± 0.1%, 99.4 ± 0.8%, 97.4 ± 2.1%, 99.5 ± 0.8%, 91.5 ± 9.3%, 96.2 ± 1.6%; *n* = 6 rats; see also **Figures 1C** and **S5G**).

#### Place-field directionality

For each place cell, a directionality index (DI) was calculated based on firing rates in the preferred (*FR_pref_*) and non-preferred (*FR_npref_*) running directions of the left or right trajectories (**Figure 1G**) as (*FR_pref_* – *FR_npref_*) / (*FR_pref_* + *FR_npref_*), similar to previous studies (Navratilova et al., 2012; Ravassard et al., 2013). A directionality index of 0 indicates identical firing in both directions, whereas 1 indicates firing in one direction only. The similarity of the place-field population in two running directions was computed using the population vector overlap (PVO; **Figures 1H** and **6F**) (Ravassard et al., 2013). The population vector (PV) was the activity vector of all place cells in a certain linear position bin. The PVO was defined as the vector dot product between the PVs across all linear positions in two running directions:

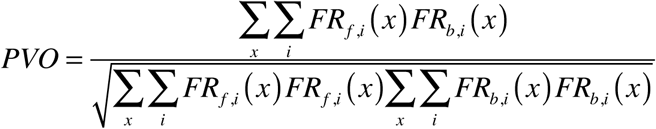

where *FR_f,i_*(*x*) is the firing rate of the *i*-th place cell at the linear position *x* along the track in a forward running direction, and *FR_b,i_*(*x*) is for the backward running direction. The PVO ranges from 0 to 1, with 1 representing identical population place-field templates in two running directions. To determine the significance values for the PVO and DI, we created 1,000 shuffle surrogates by randomly assigning a running direction to the spikes of a given cell that occurred on a given side of the maze (left or right), and computed PVO and DI from the shuffled data. Unidirectional cells were defined as cells with a DI significantly higher than its shuffle surrogates (*p* < 0.05; See **Table S1** for the number of unidirectional cells of each animal).

#### Replay decoding

Replay decoding was implemented as previously described (Davidson et al., 2009; Karlsson and Frank, 2009; Tang et al., 2017). Candidate events were defined as the SWR events during which ≥ 5 place cells fired. To detect replay, each candidate event was divided into 10-ms non-overlapping bins, and a memoryless Bayesian decoder was built for each of the four trajectory types to estimate the probability of animals’ position given the observed spikes (Bayesian reconstruction; or posterior probability matrix): *P*(*X*| **spikes**) = *P*(**spikes**| *X*)*P*(*X*)/*P*(**spikes**), where *X* is the set of all linear positions on the track of the given trajectory type, and we assumed a uniform prior probability of *X*. Assuming that all *N* place cells active during a candidate event fired independently and followed a Poisson process:

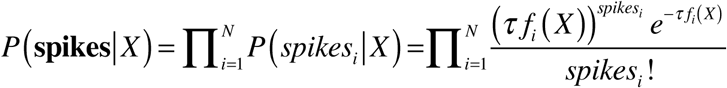

where *τ* is the duration of the time window (i.e., 10 ms for replay events, and 500 ms for active behavior), *f_i_*(*X*) is the expected firing rate of the *i*-th cell as a function of sampled location *X*, and *spikes_i_* is the number of spikes of the *i*-th cell in a given time window. Therefore, the posterior probability matrix can be derived as follows:

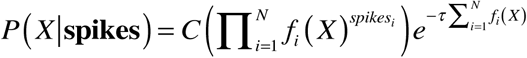

where *C* is a normalization constant. The assessment of replay events for significance was implemented as previously described (Karlsson and Frank, 2009). The *p*-value was calculated based on a Monte Carlo shuffle. First, we drew 10,000 random samples from the posterior probability matrix for each decoded bin and assigned the sampled locations to that bin. Then, we performed a linear regression on the bin number versus the location points. The resulting *R*-squared was compared with 1,500 regressions, in which the order of the temporal bins was shuffled (i.e., time shuffle) (Foster, 2017; Trimper et al., 2017). A candidate event with *p* < 0.05 based on the time shuffle was considered as a replay event. As mentioned above, each of the four trajectory types was independently decoded, and the replay trajectory was determined as the one with the lowest *p*-value from the shuffling procedure (Tang et al., 2017), of which the *R*-squared (or *r*) was reported as replay quality. Since there was a bias towards reward locations for place cells and their associated replay events (**Figures S5G** and **S5I**), similar to previous reports (Dupret et al., 2010; Pfeiffer and Foster, 2013), we excluded the spatial positions within 15 cm of reward wells from the place-field templates to detect replay (all our main results were similar without the 15-cm exclusion) to ensure that this bias did not affect our detection of the replay events representing an animal’s moving path. For plotting purposes only, a moving window (20 ms advanced in steps of 10 ms) was used for displaying replay sequences (**Figures 2**, **3** and **S3**) (Farooq and Dragoi, 2019). Only behavioral sessions with more than one replay event per analyzed category were included for calculating the percentage (**Figures 3J-3L**, **Figures 5E** and **5F**).

#### Replay prediction

For replay prediction (**Figures 3M** and **3N**, **Figures 5G** and **5H**), trial-by-trial classification analysis was performed using support vector machines (SVMs) through the libsvm library (version 3.12) (Chang and Lin, 2011). During immobility periods at a given reward well (see *Behavioral state definition*), the number of each replay event type was used as a feature (*n* = 8 possible features, 4 trajectory types × 2 replay orders, i.e., forward and reverse). Unless otherwise noted, all classifiers were *C*-SVMs with a radial basis function (Gaussian) kernel and trained on behaviorally correct trials. Hyperparameter (*C* and *γ*; regularization weight and radial basis function width, respectively) selection was performed using a random search method with leave-one-out cross-validation to prevent overfitting. The selected hyperparameters were then used to report the leave-one-out cross-validation accuracy. The percentage of correctly inferred trials was computed across all training/test trial combinations to give prediction accuracy. The significance of this prediction was determined by comparing to the distribution of shuffled data. Each “shuffled” dataset was constructed by randomly shuffling the trial labels (see below), and this shuffled dataset was used to train a classifier in the same way as the actual dataset. A prediction accuracy based on the actual dataset that was higher than the shuffled ones with *p* < 0.05 was considered as significant.

Specifically, to classify well identity (**Figures 3M** and **3N**), two independent SVMs were trained on forward and reverse replay, respectively. For a given replay order (i.e., forward or reverse), the number of each replay event type during immobility at a given well was used as a feature (*n* = 4 features; 4 trajectory types) and the well ID was used as the trial label (*k* = 3; center, right and left wells). For this prediction, a trial (or transition) is therefore defined based on the immobility period at the well during a given behavioral sequence. Only transitions where at least one replay event occurred for a given replay order were used. Because the incorrect trials mostly occurred during learning of the outbound rule (**Figure S2**) (Jadhav et al., 2012), these incorrect trials were selected to compare the replay predictions of correct vs. incorrect choices. During these incorrect trials, the animal was at the center well and performed a correct past trajectory (inbound), but was about to choose the next choice incorrectly (outbound). The numbers of the 4 replay event types during immobility at the center well for these incorrect trials were used as input features (*n* = 4 features) to predict the well IDs (*k* = 3; center, right and left wells) using the SVMs trained on all correct trials. The percentage of these trials that correctly predicted the center well was reported as prediction accuracy. To calculate statistical significance, correct trials at the center well were randomly subsampled 1,000 times to match the number of incorrect trials for computing prediction accuracy (**Figure 3N**).

To predict actual taken vs. not-taken paths based on replay (**Figures 5G** and **5H**), independent SVMs were trained for each learning stage (i.e., early, middle and late) and replay order (*n* = 6 SVMs, 3 learning stages × 2 replay orders) at either center or side wells. For a given replay order (i.e., forward or reverse), the numbers of events replayed for all 4 possible paths during the immobility period at a given well were used as features (*n* = 4 features), and the taken behavioral sequence was used as the trial label (*k* = 2; taken vs. not-taken sequences; see **Figure 1B**). Only correct trials with at least one replay event in the given replay order were used for prediction.

#### CA1-PFC population reactivation analysis

In substance, the method to measure CA1-PFC reactivation here is similar to the “template matching” or “reactivation strength” approaches used in several previous studies (Euston et al., 2007; Girardeau et al., 2017; Kudrimoti et al., 1999; Lansink et al., 2009; Peyrache et al., 2009; Tang et al., 2017; Wilber et al., 2017), but operated on a timescale of replay dynamics to examine finer temporal structure of the reactivation activity (i.e., 10 ms instead of 50-100 ms binning in the previous studies). Reactivation candidate events were defined as the SWR events during which ≥ 5 place cells and ≥ 5 PFC cells fired; therefore, they represent a subset of replay candidate events that were defined using only the CA1 place cell criterion. For a candidate event with *N* CA1 and *M* PFC cells firing (*N* ≥ 5, and *M* ≥ 5), a (*N* × *M*) synchronization matrix during RUN (C_RUN_) was calculated with each element (*C_i,j_*) representing the Pearson correlation coefficient (*C_i,j_*) of the linearized spatial maps on a certain trajectory type (2-cm bin) of the *i*-th CA1 cell and the *j*-th PFC cell. To measure the population synchronization pattern during the SWR, the spike trains during the candidate event were divided into 10-ms bins as in the CA1 replay analysis and *z* transformed:

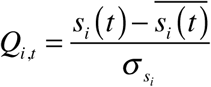

where *s_i_*(*t*) is the spike train of the *i*-th cell during the candidate event, and 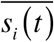 and *σ_si_* are the mean and standard deviation of *s_i_*(*t*), respectively. The (*N* × *M*) synchronization matrix during the candidate event (C_SWR_) was then calculated with each element (*C_i,j_*) representing the correlation of a CA1-PFC cell pair: 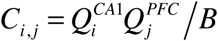, where *i* ≤ *N*, *j* ≤ *M*, and *B* is the total number of time bins during the SWR. The reactivation strength of this event was measured as the correlation coefficient (*R*) between the population matrices, C_RUN_ and C_SWR_. To evaluate the significance of the reactivation strength, the spike times during the SWR were randomly shuffled 1,500 times, in order to randomize the synchronization between CA1 and PFC cells, but conserve the structure of their spatial maps. A candidate event with *p* < 0.05 versus its shuffled data was considered as a reactivation event. As in the replay analysis, the reactivated trajectory was determined as the one (among the four possible trajectory types) with the lowest *p*-value determined by the shuffling procedure. This was used to identify coherent CA1-PFC ensemble reactivation that was aligned with CA1 replay, and a graphical illustration of the method is provided in **Figure S8**. We used this synchronization measure because a synchronous, rather than sequential, timing relationship of cross-regional reactivation, including CA1-PFC reactivation, was reported in previous studies (Girardeau et al., 2017; Lansink et al., 2009; Tang et al., 2017), and it also allows us to directly compute and examine the combined cross-regional reactivation, rather than detecting reactivation separately in each region and then measuring their correlation.

#### Model simulations for reactivation analysis

We created a simulated neuronal population as an illustrative example of the reactivation method described above (**Figure S8**), in comparison to other potential methods. We used model simulations here because while the “true” connectivity of recorded CA1-PFC populations is inaccessible, using the forward-modelling scheme, in which the “ground truth” is known and the neuronal connectivity among the simulated population can be defined, allows for validation of the method for identifying synchronous reactivation. For simplicity, we formulate the model for 5 CA1 and 5 PFC cells. From neurophysiological data, it is known that when an animal moves along a trajectory, CA1 place cells often exhibit narrowly tuned single-peaked place fields, whereas the spatial maps of PFC cells are often broadly tuned and multi-peaked, suggesting a many-to-one mapping between hippocampal and PFC representations (Jadhav et al., 2016; Tang et al., 2017; Yu et al., 2018). Motivated by these response properties, the estimated firing rate (i.e., place field) of each place cell *i*, *r_i_* (*t*), is defined as a Gaussian tuning curve,

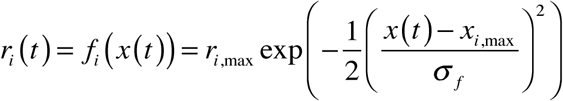

where *r_i_*_,max_ is the maximum firing rate of the *i*-th cell, *x* (*t*) is the linear position of the animal at time *t*, *x*_*i*,max_ is the position evoking the maximum average rate *r_i_*_,max_ of the cell, and *σ_f_* determines the width of the tuning curve (*σ_f_* = 5 cm for CA1 cells). The synchronization pattern between CA1 and PFC cells within the population is defined by a many-to-one connectivity matrix. If the activity of the *k*-th PFC cell is synchronized with that of *n* CA1 cells (*n* ≥ 1), the estimated firing rate (i.e., spatial map) of the *k*-th PFC cell, *r_k_* (*t*) is determined as,

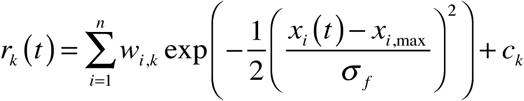

where *w_i_*_,*k*_ is the connectivity weight between the *i*-th CA1 cell (*i* = 1, 2, …, *n*) and the *k*-th PFC cell, *c_k_* is the baseline firing rate of the *k*-th PFC cell, and *σ_f_* is 10 cm for PFC cells. Thus, the peak positions of the PFC spatial map were determined by the synchronized CA1 cells as {*x*_*i*,max_ } for *i* = 1, 2, …, *n*. We assume that the spiking activity of each neuron follows an inhomogeneous Poisson process. Thus, spike sequences were simulated by using the estimated firing rate *r* (*t*) to drive a Poisson process. The probability of observing *N_i,t_* spikes of the *i*-th cell in a bin of size *τ* is given by a Poisson distribution with a rate parameter *r_i_* (*t*)*τ*,

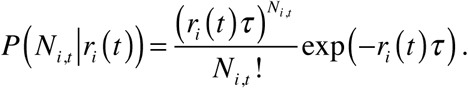

As in the real data, *τ* is 10 ms for SWR events and 500 ms for active behavior. Note that this Poisson process generates an irregular firing pattern during SWRs that reflects the underlying spatial map. The reactivation method was then applied to the simulated data as described above, and compared to Bayesian decoding (O’Neill et al., 2017) and line-fitting (Ólafsdóttir et al., 2016, 2017) methods.

#### CA1-PFC activation ratio and pairwise reactivation

For each cell activated during a SWR synchronous event (i.e., candidate event), the average firing rate of the cell during the SWR divided by its average firing rate across the behavioral session was used as its activation ratio. The activation ratio for a SWR synchronous event was then measured as the mean activation ratio across all cells activated during the SWR (**Figure 7H**, **Figure S7E**). Pairwise reactivation strength was measured as the correlation coefficient between spatial correlation and SWR cofiring of cell pairs (**Figure 7I**, **Figure S7G**), as described previously (Tang et al., 2017). In brief, the spatial correlation of a cell pair was defined as the Pearson’s correlation coefficient between their linearized spatial maps across all 4 trajectory types. SWR cofiring of a cell pair was calculated as the Pearson’s correlation coefficient between their spike trains occurring during SWR events using 50-ms bins.

#### Statistical analysis

Data analysis was performed using custom routines in Matlab (MathWorks, Natick, MA). We used nonparametric and two-tailed tests for statistical comparisons throughout the paper, unless otherwise noted. We used repeated measures ANOVA for multiple comparisons of paired Gaussian distributions, followed by a Tukey’s test, when appropriate. For non-Gaussian distributions of multiple groups, we used Kruskal-Wallis or Friedman test, with *post hoc* analysis performed using a Dunn’s test. *P* < 0.05 was considered the cutoff for statistical significance. Unless otherwise noted, values and error bars in the text denote means ± SEM.

### DATA AND SOFTWARE AVAILABILITY

The data and code that support findings of this study are available upon reasonable request by contacting the Lead Contact, Dr. Shantanu P. Jadhav.

## Supplemental Information

**Figure S1.**
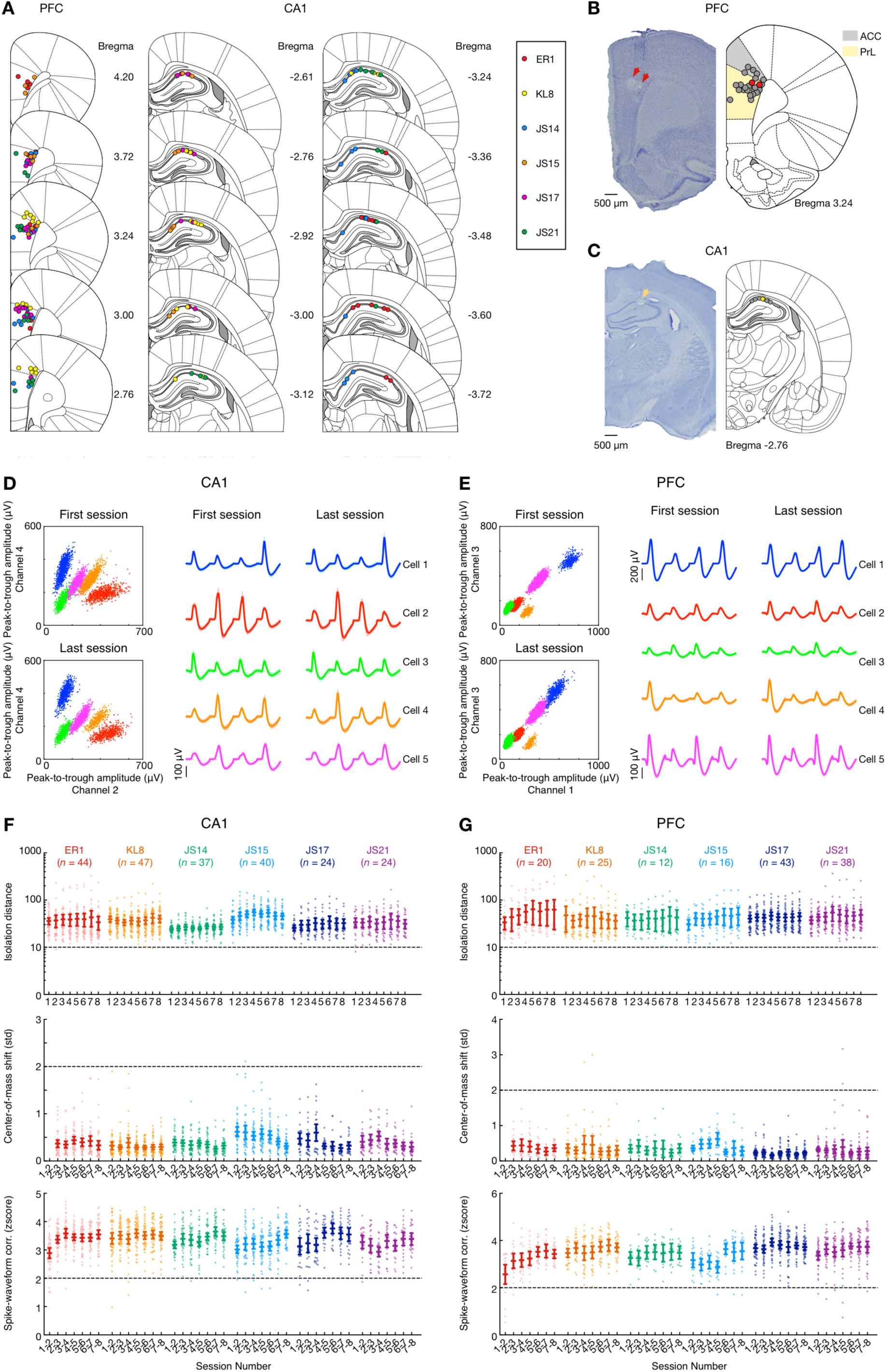
(Related to Figure 1). Recording locations, cluster quality and stability of neural recordings. **(A)** Recording locations for all 6 rats (color coded). The recording sites were reconstructed from the electrolytic lesions in *post hoc* Nissl-stained coronal brain sections and mapped onto the stereotaxic atlas (Paxinos and Watson, 2004). The distance from Bregma (mm) is denoted on the right for each section. Electrodes were localized to target areas in dorsal area CA1 and medial prefrontal cortex (PFC) after histology (PFC electrodes primarily in PreLimbic (PrL) cortex, with a few electrodes from one animal KL8 in Anterior Cingulate Cortex (ACC) area of PFC). **(B** and **C)** Representative histological sections through **(B)** PFC and **(C)** dorsal hippocampus. *Left*: Coronal Nissl-stained section illustrating lesion locations at end of tetrode tracks (marked by arrowheads). *Right*: Schematic illustration of the section shown on the left based on the stereotaxic atlas (Paxinos and Watson, 2004) (colored dots represent the lesion locations on the section). **(D** and **E)** Cluster examples of 5 single cells (color-coded) recorded on **(D)** one CA1 and **(E)** one PFC tetrode from the first and last behavioral sessions, respectively. Scatter plots show the peak-to-trough amplitudes of spike waveforms recorded on 2 of the 4 channels (each dot representing a single sampled spike; spikes associated with each isolated single unit are shown in a different color). Spike waveforms of the example cells on 4 channels are shown on the corresponding right panels (line: mean; shading: SD). **(F** and **G)** Cluster quality for all animals across 8 behavior sessions, for **(F)** CA1 place cells and **(G)** PFC cells. For the W-maze single-day learning paradigm, single-units were tracked continuously across interleaved run and rest sessions (W-maze run sessions: 17.9 ± 1.0 mins per session, 8 sessions per rat; interleaved rest sessions in rest box: 23.0 ± 4.9 mins per session, 9 sessions per rat; total recording duration: 6.04 ± 0.37 hours per rat; mean ± SD). From top to bottom, isolation distance (Schmitzer-Torbert et al., 2005), cluster center-of-mass shift (Mallory et al., 2018) across sessions (spikes from the first session of the Animal ER1 were clustered separately and therefore excluded from the center-of-mass shift analysis), correlation coefficient of spike waveforms in two consecutive sessions (Li et al., 2017) (Fisher-transformed for normality). Each dot on the scatter plots represents an isolated single unit (All cells that were continuously tracked across all 8 behavioral sessions with stable spiking waveforms, and with firing fields on the W-track, are shown; CA1: *n* = 216 cells; PFC, *n* = 154 cells). Dotted horizontal lines indicate inclusion thresholds used for each criterion. Error bars: mean ± 95% CI. Animal IDs and total number of cells recorded are denoted and color-coded on the top.

**Figure S2.**
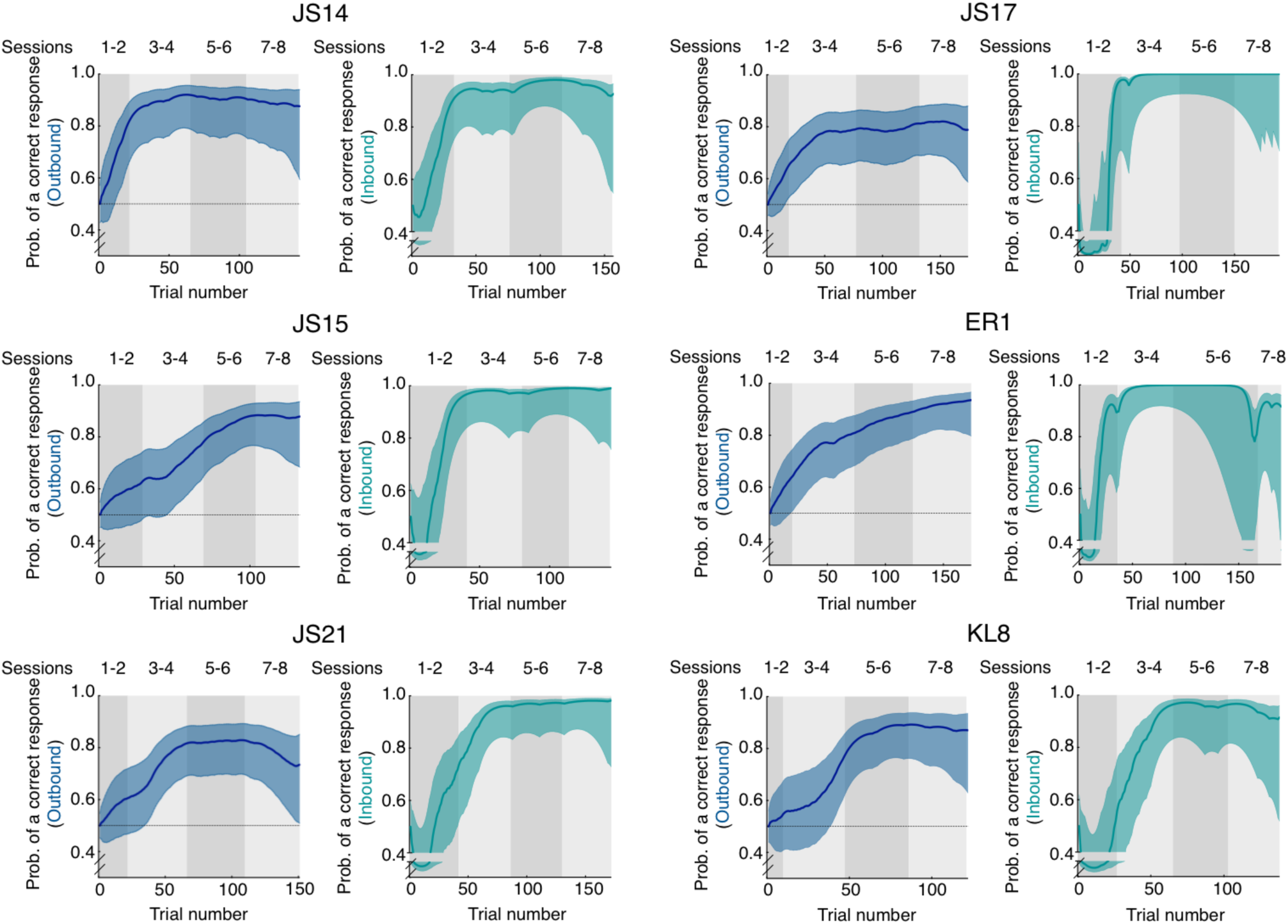
(Related to Figure 1). Behavioral performance of all animals. Data are presented as mean ± 90% CIs for the outbound (blue) and inbound (teal) components from all the 6 animals that learned the W-maze task over 8 sessions in a single-day. Learning curves were estimated using a state-space model (Jadhav et al., 2012; Maharjan et al., 2018; Smith et al., 2004). Horizontal dashed lines, chance-level performance of 0.5. Performance in first and final sessions: 53.67 ± 0.02%, and 85.17 ± 0.06% for outbound component; 33.74 ± 15.40%, and 95.45 ± 0.04% for inbound component (data are presented as mean ± SD).

**Figure S3.**
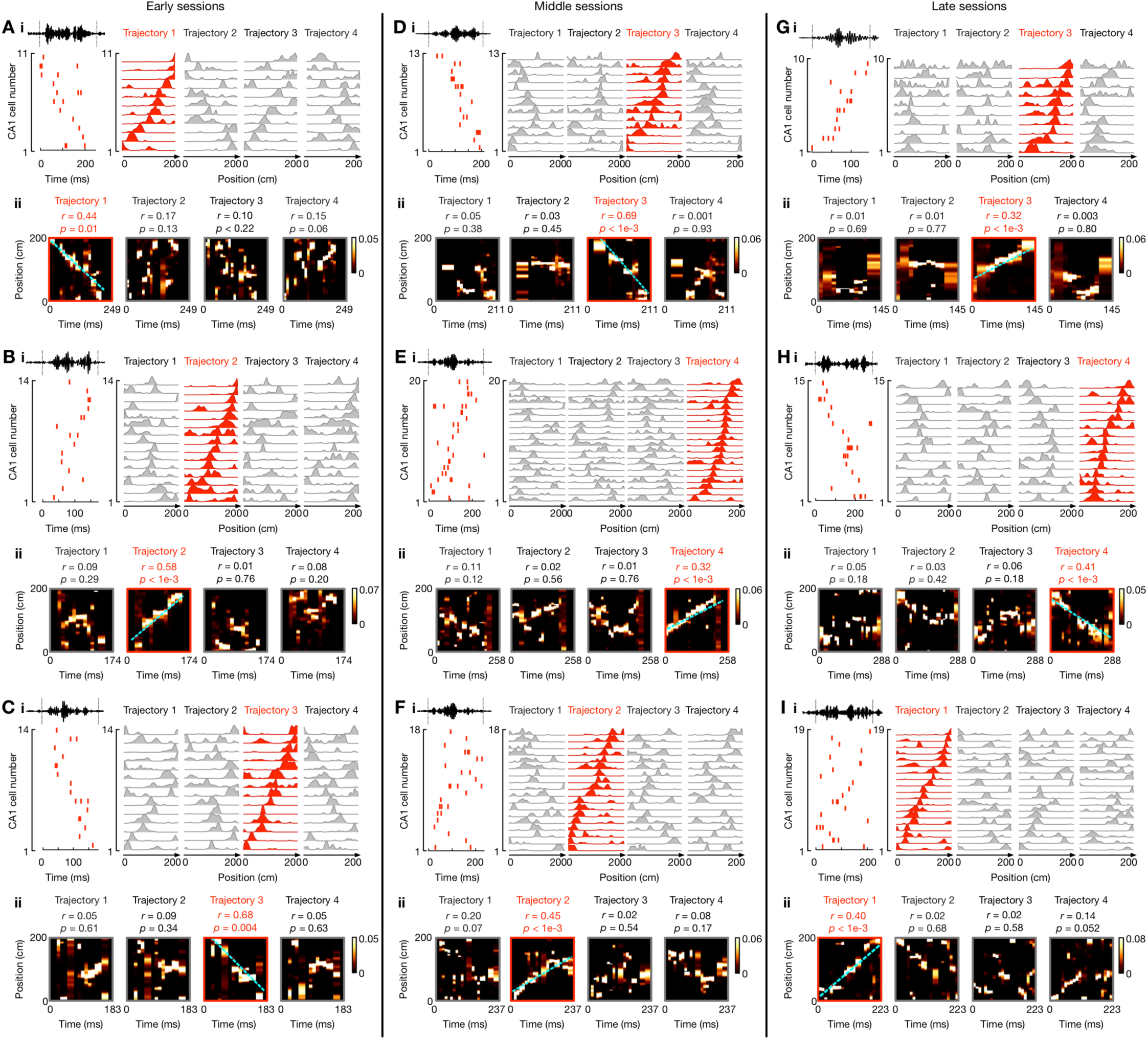
(Related to Figure 2). Additional example forward and reverse replay events of behavioral trajectories from different learning sessions. **(A-C)** Early session (Epochs 1-3) examples. **(D**-**F)** Middle session (Epochs 4-5) examples. **(G-I)** Late session (Epochs 6-8) examples. Data are presented as in Figure 2. (i) *Left*: Place-cell activity during the SWR with ripple-filtered (150-250 Hz) LFP from one tetrode shown on top (black line). *Right*: Corresponding linearized place fields on trajectories 1 to 4 sorted by their peak locations on the replay trajectory (red). **(ii)** Bayesian reconstruction of the decoded behavioral trajectories with the replay quality (*r*) and *p*-value based on time-shuffled data denoted on top. Cyan lines: the linear fit maximizing the likelihood along the replayed trajectory. Color bars: posterior probability.

**Figure S4.**
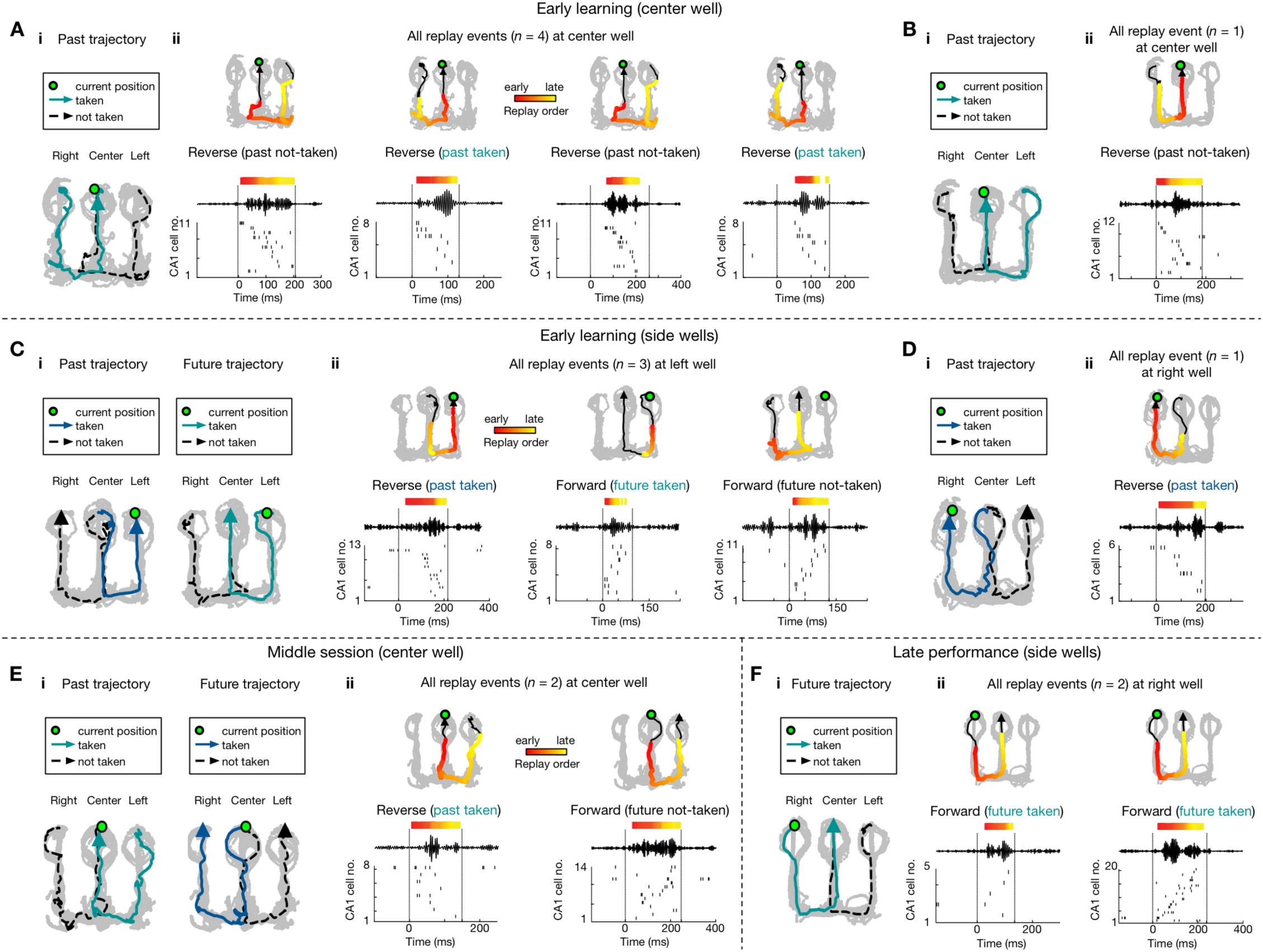
(Related to Figure 3). Representative replay events at reward wells. **(A** and **B)** Example replay events at center well during early learning (sessions 1-3). **(C** and **D)** Example replay events at side wells during early learning. **(E)** Example replay events at center well during middle sessions (sessions 4-5). **(F)** Example replay events at the side well during late performance (sessions 6-8). Data are presented as in **Figures 3G-3I**. For each example, the behavioral sequence is shown in **(i)** and all the replay events during immobility at the well (i.e., current position; green circle) are shown in **(ii)**. *Top*: Bayesian reconstruction of the replayed trajectory. Colored points overlaid on the replayed trajectory (black arrowhead line) indicate the Bayesian-decoded positions, with the color denoting relative time within the replay event (early in red, and late in yellow). *Bottom*: ripple-filtered LFP and the sequential spiking of place cells during the SWR.

**Figure S5.**
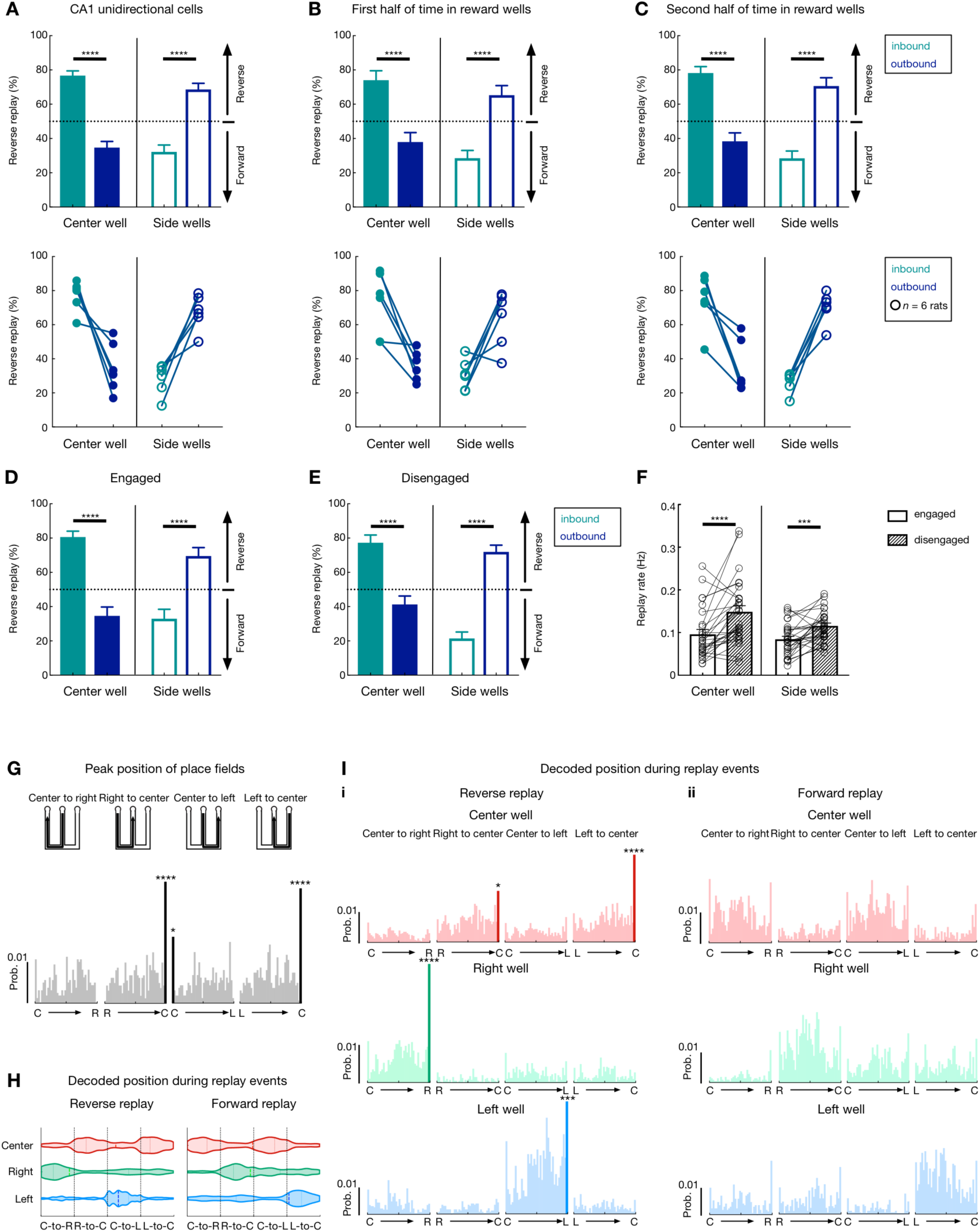
(Related to Figure 3). Control analyses examining the bias in forward and reverse replay of future and past choices. **(A)** The bias in reverse and forward replay of past and future choices is not affected by exclusion of bidirectional cells. Forward replay showed a significant bias for future choices, while reverse replay is biased for past choices, using unidirectional place cells only. Data are presented as in Figures 3J and 3K. Unidirectional cells were defined as cells with a directionality index significantly higher than the shuffle surrogates in a given session (by randomly shuffling the labels of two running directions). See **Table S1** for the number of unidirectional cells in each session. **(B** and **C)** The bias in replay content for both reverse and forward replay was seen during the first (i.e., arrival; **B**) as well as the second (i.e., departure; **C**) half of time spent in reward wells. **(D-F)** The bias in replay content did not differ appreciably during engaged **(D)** vs. disengaged **(E)** periods (Ólafsdóttir et al., 2017), although **(F)** the replay rate was significantly higher during disengaged than engaged periods. For trials with an immobility duration (defined as ≤ 4 cm/s) at a reward well longer than 6 s, the total time spent at the well was equally divided into 4 parts; the first and last parts were defined as engaged periods, and the rest were considered as disengaged. In **(F)**, each dot pair represents a session from one animal. \*\*\*\**p* < 1e-4, \*\*\**p* < 0.001, rank-sum paired tests. Error bars: SEM. **(G)** Spatial distributions of the peak positions of place fields for the 4 trajectory types (schematic of these trajectories shown on top). The *x*-axis shows the linearized positions of each trajectory with the start and end well IDs denoted (C: center well; L: left well; R: right well). Consistent with previous studies (Dupret et al., 2010; Pfeiffer and Foster, 2013), we found an accumulation of place fields at the center well (goal location; Grubb’s test for outliers, \*\*\*\**p* < 1e-4, \**p* < 0.05). Prob = probability. **(H** and **I)** Spatial distributions of the decoded positions during replay at the 3 different reward wells (color-coded). Violin plots are shown in **(H)**, and histograms are shown in **(I)**. Note that consistent with the results in Figure 3, reverse replay is biased to past paths, whereas forward replay is biased to future paths. For example, at the center well, reverse replay is biased towards positions along the past trajectories of R-to-C and L-to-C (Hartigan’s dip test for bimodality, dip = 0.062, *p* < 1e-4); whereas forward replay is biased towards positions along the future trajectories of C-to-R and C-to-L (Hartigan’s dip test for bimodality, dip = 0.065, *p* < 1e-4). In addition, consistent with previous studies (Davidson et al., 2009; Diba and Buzsáki, 2007; Foster and Wilson, 2006; Karlsson and Frank, 2009), we found that replay trajectories, specifically *only reverse replay* trajectories, tended to start from the animals’ current location at the corresponding reward well (current location is the only statistical outlier during replay at a given well; Grubb’s test for outliers, \*\*\*\**p* < 1e-4, \*\*\**p* < 0.001, \**p* < 0.05; *p* > 0.05 for forward replay shown in **I_ii_**). Note that the over-representation of current location cannot be explained solely by the spatial distributions of place fields shown in **(G)**.

**Figure S6.**
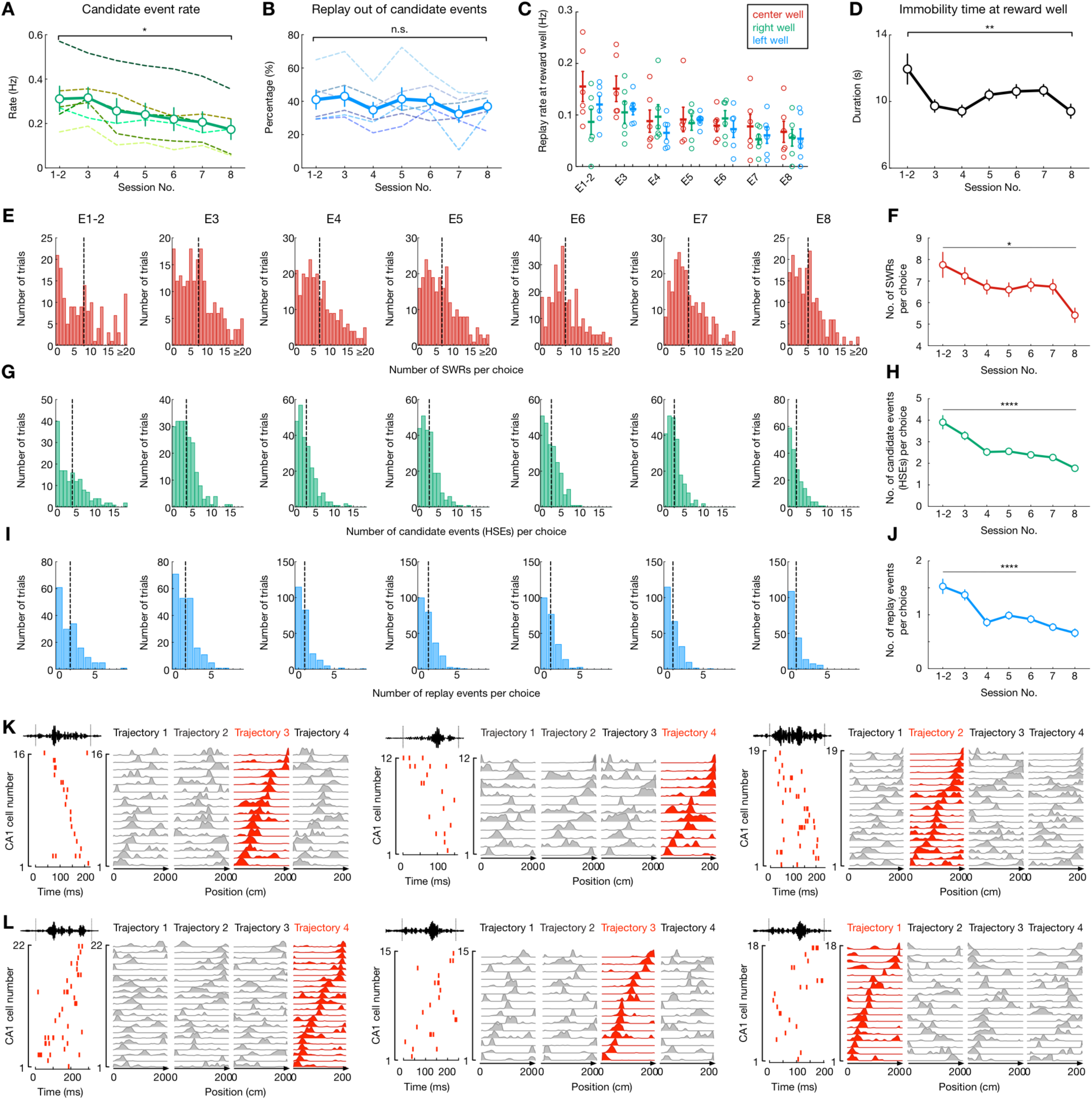
(Related to Figures 4 and 5). Change in SWR replay properties over the course of learning. **(A)** Candidate event rate significantly decreased across sessions (*p* < 0.0001 and *F*(6, 30) = 19.98, Repeated measures ANOVA; \**p* = 0.022, Tukey’s *post hoc* tests). Note that SWR rate and place-cell spiking during SWRs significantly decreased over learning, as shown in **Figure 4**. **(B)** Percentage of replay out of candidate events did not change across sessions (*p* = 0.12 and *F*(6, 30) = 2.37, repeated measures ANOVA). In **(A** and **B)**, thin dashed lines are for individual animals, thick lines and error bars represent means and SEM. **(C)** Replay rate was not significantly different at the three reward wells (*p* = 0.16, Friedman test), and replay decreased across sessions for both center and side wells (Center well: *p* = 0.0002, Friedman test; *p* = 0.0038 for sessions 1-2 vs. 8, Dunn’s *post hoc* tests; Side wells: *p* = 0.0008, Friedman test; *p* = 0.033 for sessions 1-2 vs. 7, *p* = 0.012 for sessions 3 vs. 8, Dunn’s *post hoc* tests). Each dot represents one session from a subject. Lines and error bars represent means and SEMs, respectively. **(D)** Immobility time at reward wells decreased with learning (*p* = 0.0018 and *F*(6, 1241) = 3.531, one-way ANOVA; \*\**p* = 0.0063, Holm-Sidak’s *post hoc* tests). **(E** and **F)** Number of SWRs per choice decreased across 8 behavioral sessions (\**p* = 0.027, Kruskal-Wallis test with Dunn’s *post hoc*). **(G** and **H)** Number of candidate events per choice decreased across 8 behavioral sessions (\*\*\*\**p* < 1e-4, Kruskal-Wallis test with Dunn’s *post hoc*). **(I** and **J)** Number of replay events per choice decreased across 8 behavioral sessions (\*\*\*\**p* < 1e-4, Kruskal-Wallis test with Dunn’s *post hoc*). In **E**, **G** and **I**, the vertical dashed lines on the histograms represent mean values. Error bars: SEM. **(K** and **L)** Place-field templates of the replay events in **Figure 5.** Sequential firing of place cells during replay events and corresponding linearized place fields on trajectories 1 to 4 sorted by their peak locations on the replay trajectory (red) are shown. **(K)** is for the replay events in **Figures 5A** and **5B**; **(L)** is for **Figures 5C** and **5D**.

**Figure S7.**
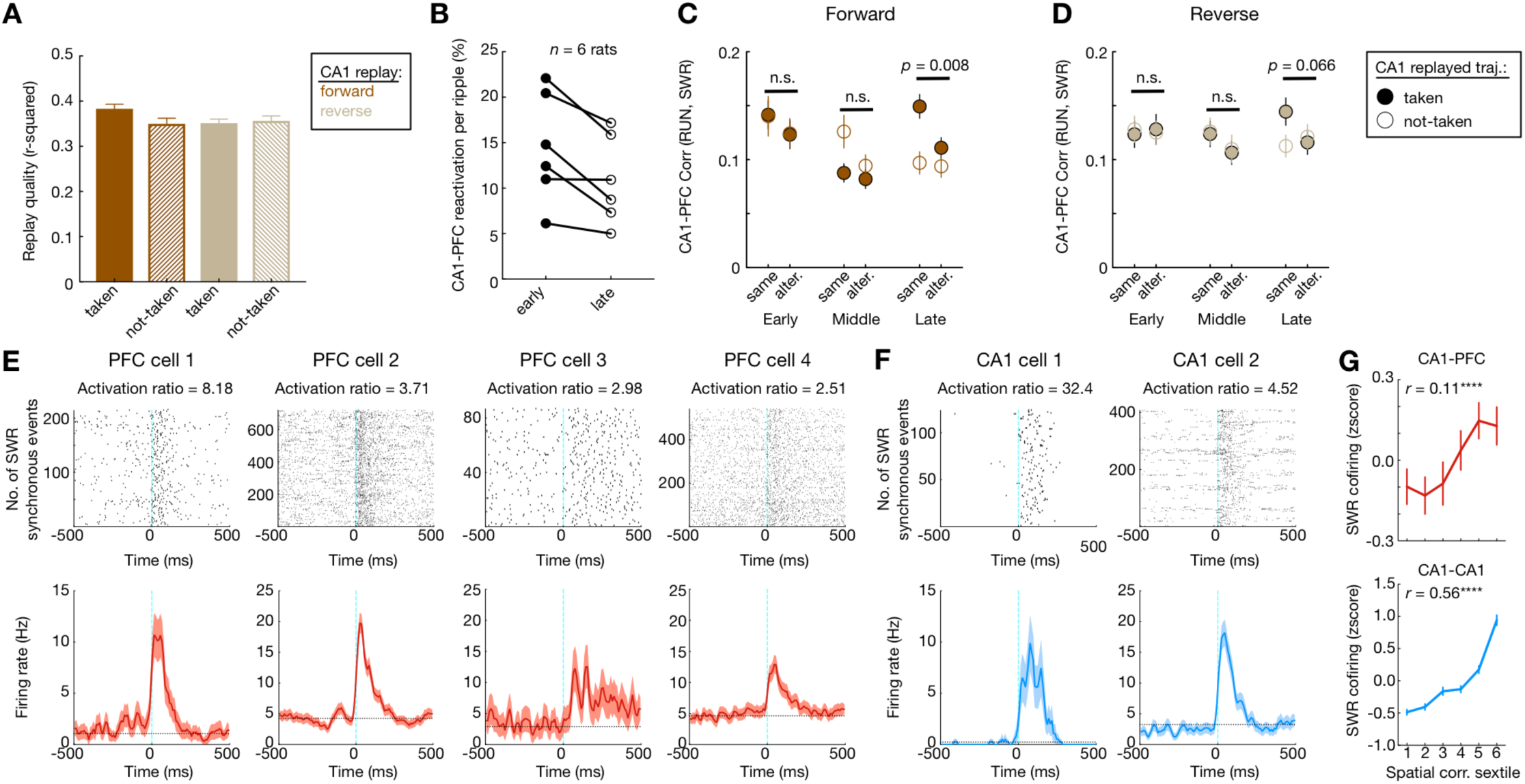
(Related to Figure 7). Properties of CA1-PFC reactivation. **(A)** CA1 replay quality of taken vs. not-taken trajectories. Replay quality was measured as “*R*-squared” from the linear regression maximizing the likelihood along the replayed trajectory. While the reactivation strength of CA1-PFC population was higher for taken vs. non-taken trajectory replay (**Figures 7E** and **7F**), we found no significant difference in CA1 replay quality for taken vs. not-taken paths (*p* = 0.066, Kruskal-Wallis test). **(B)** Number of CA1-PFC reactivation events per SWR was higher during early vs. late sessions (first 4 vs. last 4), confirming our previous reports (Tang et al., 2017). **(C** and **D)** Change of CA1-PFC reactivation strength over learning. **(C)** Stronger CA1-PFC reactivation of CA1-forward-replayed taken path compared to the not-taken path was observed only in late performance sessions (*p* = 0.21, 0.44, and 0.008 for CA1 replay of taken paths at early, middle, and late learning stages; *p* = 0.53, 0.08, and 0.66 for CA1 replay of not-taken paths at early, middle. and late learning stages; rank-sum paired tests). **(D)** No statistically significant trend was observed for reverse CA1 replay (*p* = 0.77, 0.32, and 0.066 for CA1 replay of taken paths at early, middle and late learning stages; *p* = 0.61, 0.33, and 0.57 for CA1 replay of not-taken paths at early, middle, and late learning stages; rank-sum paired tests). **(E** and **F)** Raster plots (*top*) and corresponding PSTHs (*bottom*) aligned to SWR synchronous events of four PFC **(E)** and two CA1 **(F)** neurons, showing activation during SWRs. Horizontal dashed lines: mean firing rates in the entire session. Lines and shadings in the PSTHs: means and SEMs. Activation ratio of each cell is denoted on the top of the corresponding panel. **(G)** SWR co-firing as a function of spatial correlation for CA1-PFC (*top*) and CA1-CA1 (*bottom*) cell pairs for one high-performance animal indicated in Figure 7I. Spatial correlations are divided into six subgroups with equal number of cell pairs (i.e., sextiles), similar to a previous study (Tang et al., 2017). Note the strong correlation between SWR cofiring vs. spatial correlation of CA1-PFC pairs (*r* = 0.11, *p* = 2.3e-4), as well as CA1-CA1 pairs (*r* = 0.55, *p* < 1e-10).

**Figure S8.**
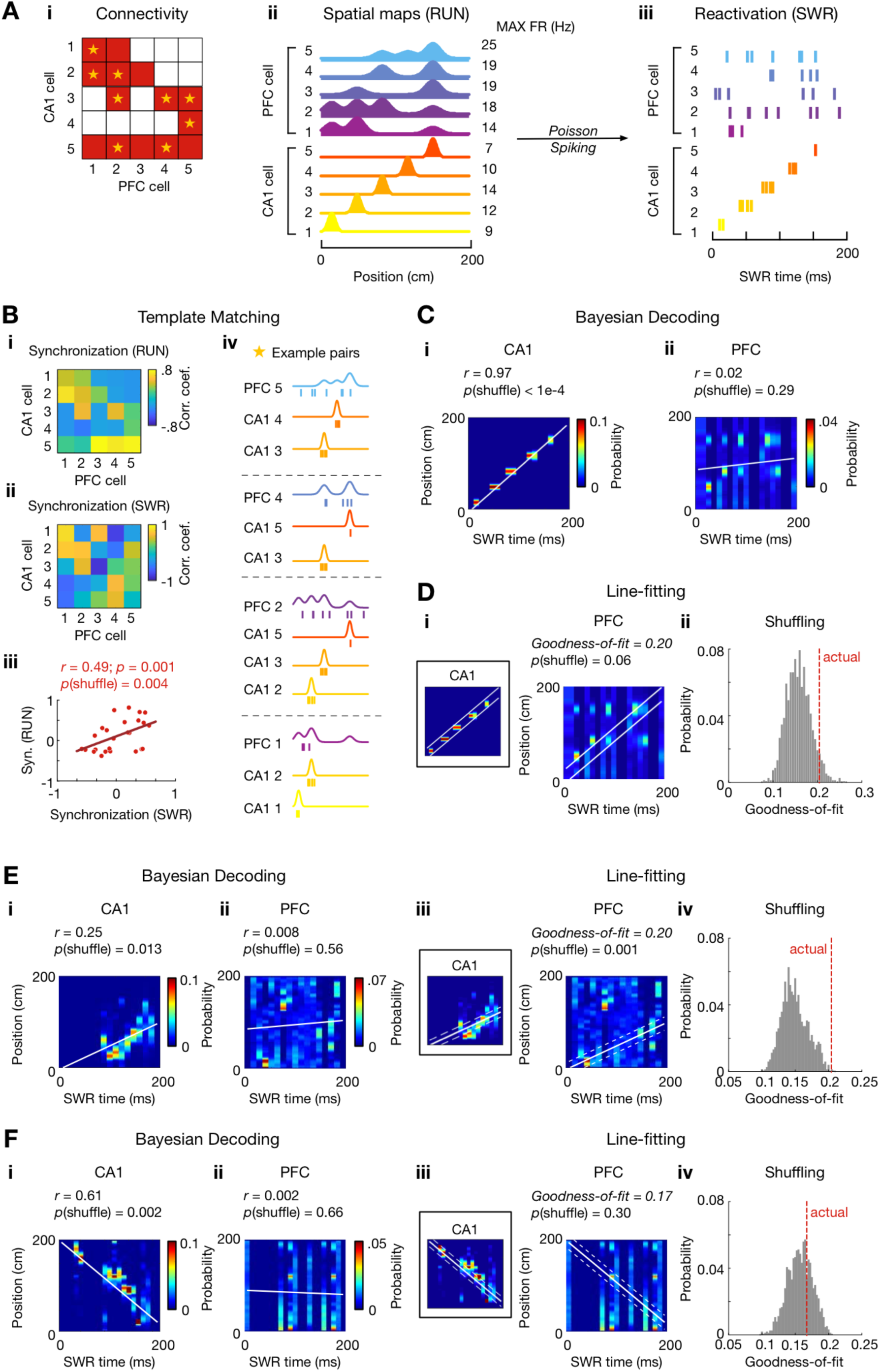
(Related to Figure 7). Template matching to detect CA1-PFC reactivation: method summary and comparison. **(A)** Generation of simulated data from a simple toy model. The simulated CA1-PFC neuronal population consists of 5 CA1 and 5 PFC cells with a many-to-one connectivity matrix. **(A_i_)** Connectivity matrix (red for CA1-PFC pairs with synchronous firing; white for non-synchronous pairs). **(A_ii_)** Spatial maps of the simulated neurons on one trajectory with their peak firing rates denoted on the right (i.e., MAX FR). PFC neurons have multi-peaked spatial maps. **(A_iii_)** Simulated spike trains during a SWR event. These spike trains were generated from a Poisson process, whose rate was determined by the spatial maps in **(A_iv_)**. **(B)** Template matching decoding of coherent CA1-PFC replay. **(B_i_)** Synchronization matrix of CA1-PFC population activity calculated during running (RUN), with each element representing the correlation of spatial maps of a CA1-PFC cell pair. **(B_ii_)** Similarly, a synchronization matrix for SWR was measured based on the pairwise correlation of spike trains during the SWR. **(B_iii_)** The reactivation strength was then computed as the correlation coefficient (*r*) between the synchronization matrices during RUN vs. SWR (each dot representing a CA1-PFC pair, red line from linear regression). For the simulated population, the reactivation strength is 0.49, significantly higher than its shuffle surrogates (randomly shuffling spike times during the SWR; *p* = 0.004; note that the SWR spike trains of the simulated data are a sample of the Poisson process during RUN). **(B_iv_)** Detailed view of the CA1-PFC coordination using example cell pairs indicated by stars on the connectivity matrix in **(A_i_)**. Curves: spatial maps. Ticks: spikes during the SWR. **(C** and **D)** Comparison of template matching, Bayesian decoding (O’Neill et al., 2017), and line-fitting (Ólafsdóttir et al., 2016, 2017) using the model in **(A)**. Despite strong correlation of the population activity during RUN and the SWR (synchronization detected using template matching in **B**), the PFC activity during the SWR was detected **(C)** as a “non-replay” event by Bayesian decoding due to a lack of sequential structure (*p* = 0.29 by comparison to time shuffles), and **(D)** not coherent with the replay in CA1 using line-fitting (*p* = 0.06 by comparison to time shuffles). **(E** and **F)** Bayesian decoding and line-fitting results of the two reactivation events shown in **Figures 7A** and **7B**, respectively. **(E)** Note that for the event in **Figure 7A** with a clear sequential structure, the line-fitting method also detected a coherent CA1-PFC reactivation (*p* = 0.001), but **(F)** not for the event in **Figure 7B** with multi-peaked, many-to-one coordination of CA1-PFC activity (*p* = 0.30 by comparison to time shuffles). White lines on Bayesian-reconstruction matrices for Bayesian decoding represent the linear fit maximizing the likelihood along the trajectory, and for line-fitting mark the extent of the line-fit based on CA1 place-cell activity (± 15 cm).

**Table S1.**
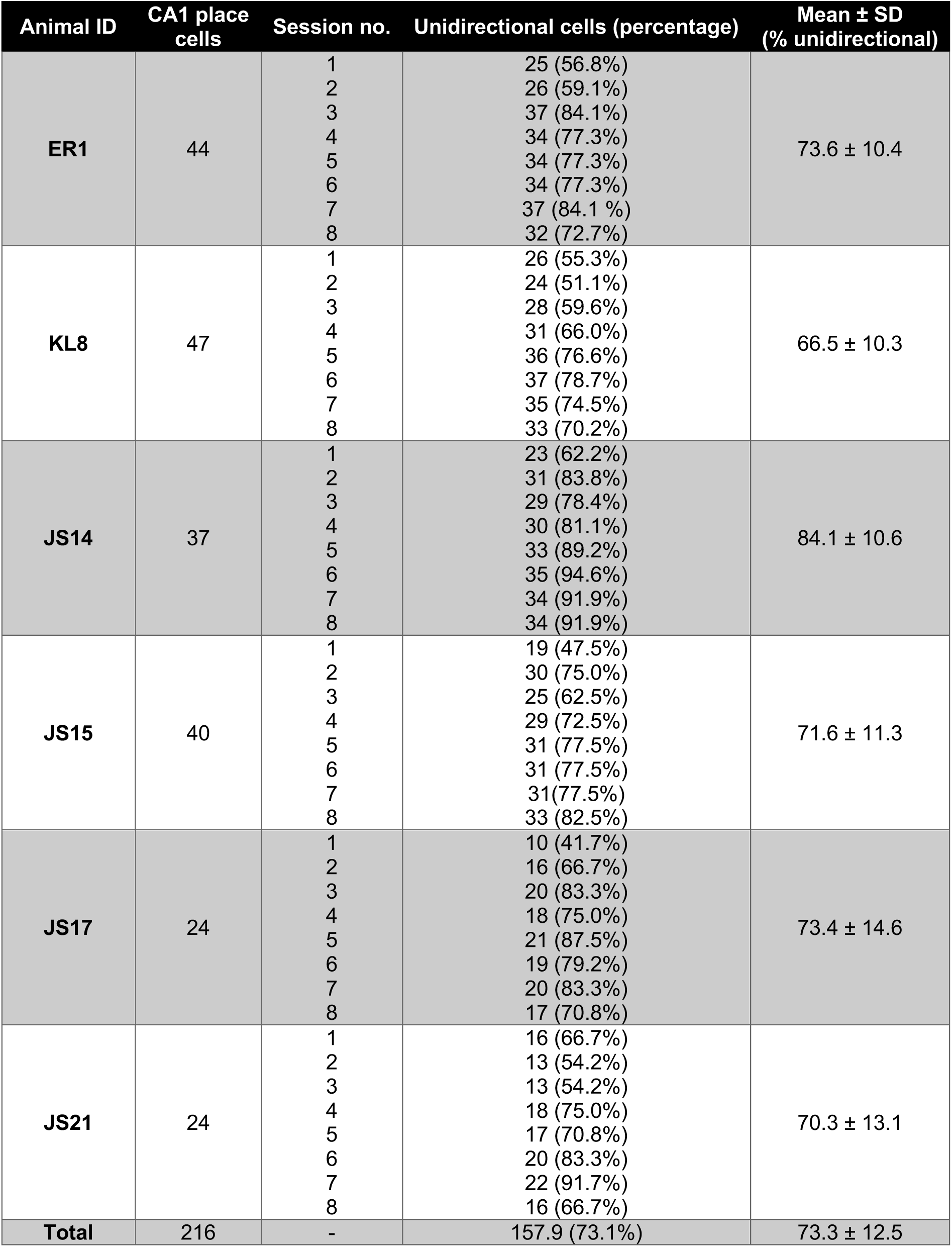
(Related to Figure 1). Summary of unidirectional cells. Number of CA1 place cells recorded continuously over 8 learning sessions, the number and percentage of unidirectional cells in each session, and percentage of unidirectional cells (mean ± SD) for each animal are shown.

| Animal ID | CA1 place cells | Session no. | Unidirectional cells (percentage) | Mean ± SD (% unidirectional) |
| --- | --- | --- | --- | --- |
| ER1 | 44 | 1 | 25 (56.8%) | 73.6 ± 10.4 |
|  |  | 2 | 26 (59.1%) |  |
|  |  | 3 | 37 (84.1%) |  |
|  |  | 4 | 34 (77.3%) |  |
|  |  | 5 | 34 (77.3%) |  |
|  |  | 6 | 34 (77.3%) |  |
|  |  | 7 | 37 (84.1 %) |  |
|  |  | 8 | 32 (72.7%) |  |
| KL8 | 47 | 1 | 26 (55.3%) | 66.5 ± 10.3 |
|  |  | 2 | 24 (51.1%) |  |
|  |  | 3 | 28 (59.6%) |  |
|  |  | 4 | 31 (66.0%) |  |
|  |  | 5 | 36 (76.6%) |  |
|  |  | 6 | 37 (78.7%) |  |
|  |  | 7 | 35 (74.5%) |  |
|  |  | 8 | 33 (70.2%) |  |
| JS14 | 37 | 1 | 23 (62.2%) | 84.1 ± 10.6 |
|  |  | 2 | 31 (83.8%) |  |
|  |  | 3 | 29 (78.4%) |  |
|  |  | 4 | 30 (81.1%) |  |
|  |  | 5 | 33 (89.2%) |  |
|  |  | 6 | 35 (94.6%) |  |
|  |  | 7 | 34 (91.9%) |  |
|  |  | 8 | 34 (91.9%) |  |
| JS15 | 40 | 1 | 19 (47.5%) | 71.6 ± 11.3 |
|  |  | 2 | 30 (75.0%) |  |
|  |  | 3 | 25 (62.5%) |  |
|  |  | 4 | 29 (72.5%) |  |
|  |  | 5 | 31 (77.5%) |  |
|  |  | 6 | 31 (77.5%) |  |
|  |  | 7 | 31(77.5%) |  |
|  |  | 8 | 33 (82.5%) |  |
| JS17 | 24 | 1 | 10 (41.7%) | 73.4 ± 14.6 |
|  |  | 2 | 16 (66.7%) |  |
|  |  | 3 | 20 (83.3%) |  |
|  |  | 4 | 18 (75.0%) |  |
|  |  | 5 | 21 (87.5%) |  |
|  |  | 6 | 19 (79.2%) |  |
|  |  | 7 | 20 (83.3%) |  |
|  |  | 8 | 17 (70.8%) |  |
| JS21 | 24 | 1 | 16 (66.7%) | 70.3 ± 13.1 |
|  |  | 2 | 13 (54.2%) |  |
|  |  | 3 | 13 (54.2%) |  |
|  |  | 4 | 18 (75.0%) |  |
|  |  | 5 | 17 (70.8%) |  |
|  |  | 6 | 20 (83.3%) |  |
|  |  | 7 | 22 (91.7%) |  |
|  |  | 8 | 16 (66.7%) |  |
| Total | 216 | - | 157.9 (73.1%) | 73.3 ± 12.5 |

